# Nuclear translocation of LINE-1 encoded ORF1p alters nuclear envelope integrity and disrupts nucleocytoplasmic transport in human neurons

**DOI:** 10.1101/2023.08.10.552479

**Authors:** Rania Znaidi, Olivia Massiani-Beaudoin, Philippe Mailly, Héloïse Monnet, The Brainbank Neuro-CEB Neuropathology Network, Rajiv L. Joshi, Julia Fuchs

## Abstract

LINE-1 retrotransposons are emerging as possible culprits in neurodegenerative diseases. However, the molecular mechanisms underlying the pathogenic role of LINE-1 and their encoded proteins ORF1p and ORF2p are still not completely understood. While the endonuclease and reverse transcriptase activities of ORF2p have been associated with DNA damage and inflammation, no pathogenic role has yet been assigned to ORF1p. Using a neuronal model of oxidative stress displaying increased LINE-1 expression, we report here that ORF1p stress-dependently translocated into the nucleus, localized to the nuclear envelope and directly interacted with nuclear import proteins, nuclear pore complex components and the inner nuclear lamina. Stress-dependent targeting of nuclear envelope components by ORF1p altered nuclear envelope integrity, disrupted nucleocytoplasmic transport and induced heterochromatin destructuration, features associated with neurodegenerative diseases and aging. Neurons of post-mortem Parkinson disease (PD) patients and non-PD affected controls expressed ORF1p and nuclear ORF1p levels correlated with altered nuclear shape in PD. Overexpression of ORF1p in neurons in the absence of stress recapitulated nuclear envelope dysfunctions and increased nuclear ORF1p levels correlated with a loss of nuclear circularity. Stress-induced nuclear alterations were restored by blocking ORF1p nuclear import or by the small molecule remodelin. This study thus reveals a retrotransposition- and ORF2p- independent pathogenic action of ORF1p at the nuclear envelope and points to ORF1p as a novel target for neuroprotection.

## Introduction

Repetitive sequences derived from transposable elements (TEs) represent 45% of the human genome, including LTR (Long Terminal Repeat) and non-LTR retroelements. Non-LTR retrotransposons of the Long Interspersed Element-1 (LINE-1) family have been the most active TEs in mammalian genomes and LINE-1-derived sequences represent almost 21% of the human genome which carries nearly 500000 LINE-1 copies ^1^. Although most of these LINE-1 copies are remnants of past retrotransposition events and are truncated or mutated, the reference human genome still carries 146 full-length LINE-1 copies (according to LINE-1Base database) which can be autonomously active and competent for retrotransposition ^2^. The full-length LINE-1 sense strand codes for two proteins, namely ORF1p and ORF2p, which are required for retrotransposition. ORF1p is an RNA binding protein and ORF2p encompasses reverse transcriptase and endonuclease activities. Since LINE-1 activation can lead to deleterious consequences such as DNA damage and genomic instability ^3–6^, senescence and inflammation ^7,8^, these TEs are repressed in most somatic cells at the transcriptional and post-transcriptional level. Failure of these repressive mechanisms can lead to the activation of LINE-1 and other TEs in the context of aging or human diseases like cancer.

Several recent studies have reported that TE activation might be linked to neurodegenerative diseases (NDs) such as Alzheimer’s disease (AD), non-AD tauopathies, Parkinson disease (PD), Amyotrophic Lateral Sclerosis (ALS) and Huntington disease (HD) ^5,9–14^. Although NDs present many common hallmarks such as pathological protein aggregation, synaptic and neuronal network dysfunction, aberrant proteostasis, cytoskeletal abnormalities, altered energy homeostasis, DNA and RNA defects, inflammation, oxidative stress and neuronal cell death ^15,16^, the genetic basis and the molecular and cellular mechanisms underlying neurodegeneration still remain elusive. In this regard, recent work from our group ^5,17^ and others ^13,14,18^ suggests that TE activation could directly contribute to neuronal dysfunctions associated with NDs. Indeed, several pathogenic features of NDs might actually be linked to TE de-repression ^19,20^. While the deleterious consequences attributed to the endonuclease and reverse transcriptase activities of LINE-1 ORF2p (i.e. DNA strand breaks, cytosolic single-stranded DNA inducing innate immune activation and inflammation) are beginning to be well unraveled, no particular pathogenic function has yet been assigned to the RNA chaperone ORF1p.

To directly address the consequences of endogenous full-length LINE-1 activation and increased LINE-1-encoded proteins in adult neurons, we used a neuronal model of acute oxidative stress displaying an increase in LINE-1 encoded ORF1p expression. Using this model, we uncovered a so far unrecognized pathogenic role of ORF1p in potentiating nuclear envelope (NE) barrier alterations. This suggests that stressed neurons are susceptible to ORF1p mediated NE perturbations leading to the amplification of NE dysfunctions which are a novel emerging feature of many NDs.

## Materiel and methods

### Cell culture

The Lund Human Mesencephalic (LUHMES) cell line is a conditionally immortalized cell line. These cells are derived from the tetracycline-controlled, v-myc-overexpressing human mesencephalic cell line MESC2.10 ^21,22^. The culturing and handling procedure of LUHMES cells were as described previously ^21^. LUHMES cells can be differentiated into mature dopaminergic neurons. The differentiation was performed following the two-step protocol previously established ^23^. Cell culture flasks, multi well plates and glass coverslips used for LUHMES culture were pre-coated by adding 50 µg/ml poly-L-ornithine (Sigma) diluted in 1X PBS (Phosphate buffered saline) for 1 h at 37°C, washed 3 times in 1X PBS, and then incubated with 1 µg/ml fibronectin (Sigma) diluted in 1X PBS for 3 h minimum at 37°C to promote cell attachment. Uncapped flasks were air-dried before cell plating.

The base cell culture medium consisted of Advanced DMEM/F12 (ThermoFisher Scientific), N2-Supplement (ThermoFisher Scientific) and 2 mM GlutaMAX (ThermoFisher Scientific, 35050061) which was supplemented by 40 ng/ml recombinant human basic fibroblast growth factor FGF (Peprotech) during proliferation, at 37°C in 100% humidified air with 5% CO2 until 70%–80% of cellular confluence is reached.

The differentiation process was initiated by the shut-down of v-myc by adding 1 µg/ml doxycycline (Sigma) in the base medium supplemented with 2 ng/ml recombinant human glial-derived neurotrophic factor GDNF (Peprotech) and 1 mM cAMP (Sigma). The pre-differentiated neurons were replated 48 h later at a defined density of 150 000 cells/well in 24-well plates or 10^6^ cells/well in 6-well plates. To confirm successful generation of post-mitotic LUHMES, expression of neuronal and dopaminergic markers was assessed throughout the differentiation protocol, which lasts a total of 5 days (Fig. 1A; Suppl. Fig. 1A). All experiments were performed on day 5 of differentiation, at which point LUHMES cells were considered to be completely mature post-mitotic neurons.

**Figure 1.**
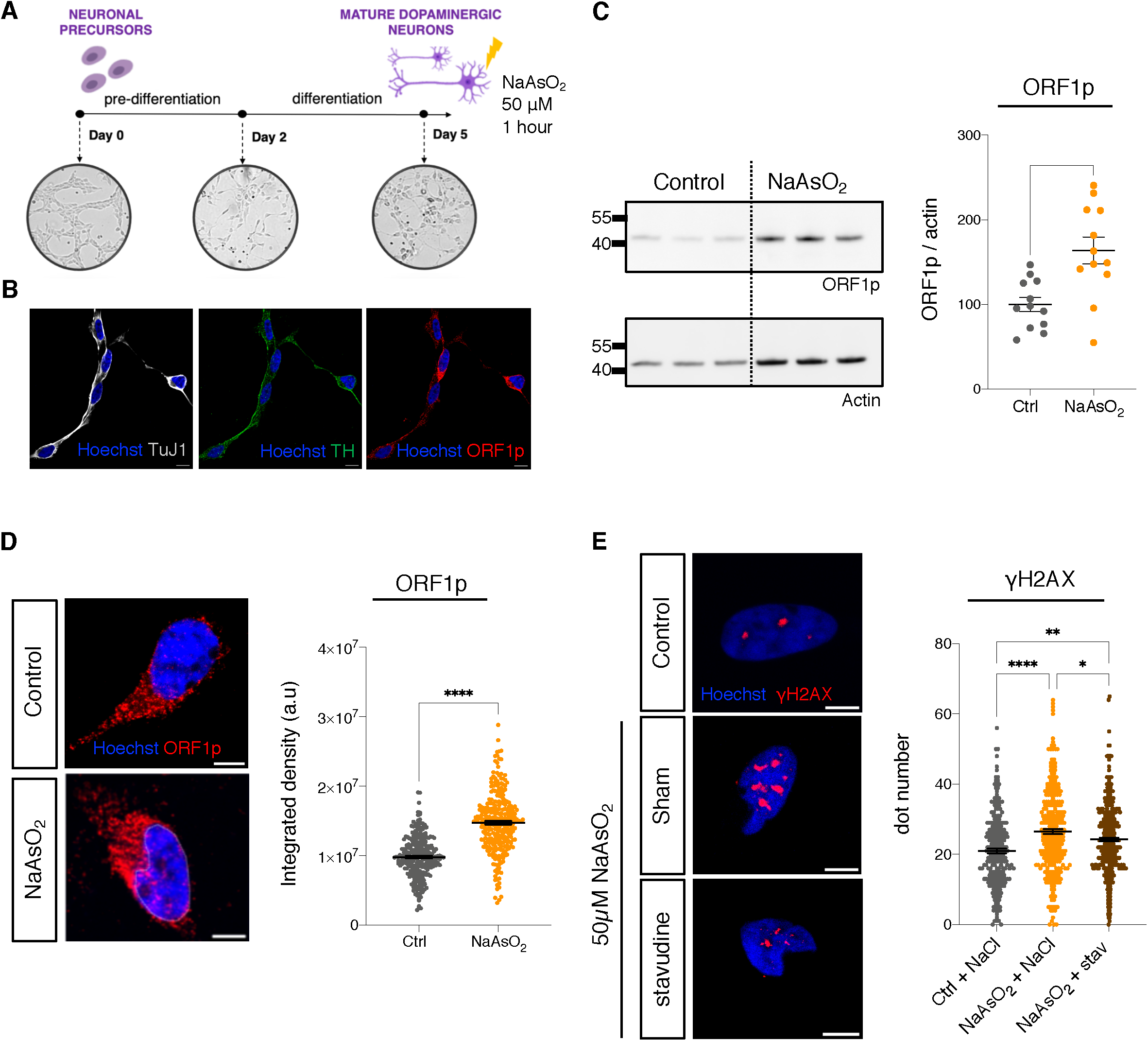
LINE-1 are upregulated in differentiated LUHMES cells following stress. (A) Experimental paradigm of the differentiation scheme of LUHMES neuronal precursor cells into mature, post-mitotic dopaminergic neurons. At day 5 of differentiation, neurons were subjected to an oxidative stress using arsenite (NaAsO_2_). (B) After 5 days of differentiation, LUHMES cells express the neuronal marker TuJ1 and the dopaminergic marker TH as revealed by immunofluorescence. These neurons express LINE-1 at steady-state as shown by immunostaining for ORF1p protein. (C) Differentiated LUHMES neurons subjected to arsenite treatment, show increased levels of ORF1p protein by Western blot analysis. n = 3 wells per condition from 4 independent experiments, mean ± SEM, two-tailed t test. (D) Immunostaining for LINE-1 ORF1p of differentiated LUHMES cells treated with arsenite and quantification of global ORF1p intensity, n > 100 neurons were quantified per condition, a representative experiment (3 wells per condition) of 3 independent experiments, mean ± SEM, two-tailed t test. (E) Immunostaining of γH2AX following arsenite treatment in the presence or the absence of stavudine and quantification of γH2AX foci number in the nucleus. n > 100 neurons were quantified per condition, a representative experiment (3 wells per condition) of 2 independent experiments, mean ± SEM, One-way ANOVA with Tukey’s multiple comparisons test (*p < 0.5, **p < 0.01, ***p < 0.001, ****p < 0.0001. (Also see supplementary Fig. 1). Scale bar, 5 µm.

### Arsenite-induced oxidative stress

Stock of sodium arsenite solution (NaAsO_2_) (Sigma) was stored at room temperature. Dilutions were carried out shortly before each experiment. Fully differentiated LUHMES cells (at day 5 of differentiation) were subjected to arsenite during 1 h at 37°C at a concentration of 50 µM to induce oxidative stress.

### Drug treatment

For treatment with remodelin (a gift from Drs. R. Rodriguez and L. Colombeau), differentiated neurons at day 4 of differentiation, were incubated in fresh culture medium containing remodelin (10 µM) or DMSO for 24 h and then subjected to oxidative stress using arsenite (50 µM) for 1 h (in the presence of remodelin). For stavudine treatment, differentiated neurons at day 4 of differentiation, were incubated in fresh culture medium containing stavudine (10 µM) or NaCl for 24 h, and then subjected to oxidative stress using arsenite (50 µM) for 1 h (in the presence of stavudine). For treatment using nuclear import inhibitors, differentiated neurons at day 5 of differentiation, were incubated in fresh culture medium containing ivermectin (5 µM; CliniSciences) and importazole (10 µM; Scientific Laboratory Supplies) or DMSO for 2 h, and were then treated with arsenite (50 µM) for 1 h. In all cases, neurons were then fixed and processed for immunocytochemistry.

### Transfection methods

Adherent electroporation (Nucleofection): LUHMES cells were seeded at a density of 150 000 cells/well into 24-well plates already containing cover-slips pre-coated with poly-L-ornithine and fibronectine and were differentiated as described in Cell culture section. After 3 days of differentiation, cell culture medium was replaced by pre-warmed fresh medium and incubated for 1 h at 37°C prior to transfection. LUHMES cells were transfected by Nucleofection using the AMAXA 4D-Nucleofector® Y Unit (Lonza) which enables the direct Nucleofection (by electroporation) of cells in adherence in 24-well culture plates using the AD1 4D-Nucleofector™ Y Kit (Lonza, V4YP-1A24) for neuronal cells electroporation. The ER-100 program was the most appropriate to LUHMES cells in terms of transfection efficiency and viability and was used for all subsequent transfections. One Nucleofection sample contained 150 000 cells/well, 10 µg plasmid DNA (pcDNA3-ORF1-HA, expressing ORF1p with an N-Ter HA tag; a gift from Dr. J. Moran and Dr. Y. Ariumi) or 10 µg pmaxGFP vector and 350 µl AD1 4D-Nucleofector Y solution. The 24-well dipping electrode array (provided in the kit) was inserted into the plate and cells were transfected by using the ER-100 Nucleofector program. After nucleofection, AD1 solution was removed and the cells were immediately washed once with fresh medium and incubated in humidified 37°C / 5% CO2 incubator. Media was not changed until analysis at day 5 of differentiation (48 h after transfection). Transfection efficiency (48 h after transfection) was monitored by fluorescence microscopy.

### Protein extraction

After aspirating media, cells were washed one time with 1X PBS. 200 to 500 µl of 1X RIPA lysis buffer (50 mM Tris-HCL pH 8, 150 mM Nacl, 0,5% Sodium deoxycholate, 1% NP-40, 0,1% SDS) with 1X protease inhibitor (Sigma) was added to each well in the plate. Cells were scraped and lysates were transferred to a sterile microtube. The lysates were incubated on ice for 30 min and sonicated using the Bioruptor UCD-200 for 15 min (30s on/30s off) on ice at high power to shear contaminating DNA. After collecting the supernatant (avoiding the pellet) into new microtubes, protein concentration was determined using Pierce^TM^ BCA Protein Assay Kit. Aliquot of lysates were stored at −20°C avoiding multiple freeze/thaw cycles. Proteins were taken in 1X Laemmli buffer (250 mM Tris pH 6,8, 10% sodium dodecyl sulfate, 1,25% bromophenol blue, 5% β-mercaptoethanol, 50% glycerol) containing DTT, boiled 10 minutes at 95°C and stored at −20°C until analyzed.

### Western blot analysis

A 1.5 mm NuPAGE 4-12% Bis-Tris-Gel (Invitrogen™) polyacrylamide gel was used for Western blots. 10 µg of protein were loaded per well. Gel migrated in NuPAGE™ MES SDS running buffer (1X) (Invitrogen™) for 55 minutes at 200 V. Gels were transferred into a methanol activated PVDF membrane (Immobilon) in a transfer buffer (Tris 25 mM, pH 8,3 and glycine 192 mM) during 1 h at 400 mA at 4°C. For blocking, 5% milk in TBST (0,2% Tween 20, 150 mM NaCI, 10 mM Tris pH:8) was used during 1 h at room temperature. The membranes were then incubated overnight with primary antibody anti-ORF1p 1:500 (Abcam, ab245249). The membranes were incubated with anti-actin peroxidase antibody 1:20000 (Sigma) diluted in 5% milk in TBST, 1 h at room temperature on a shaker. The membranes were washed in TBST (1X) and incubated with secondary antibody including anti-rabbit HRP 1:2000 (Cell Signaling) and anti-mouse HRP 1:2000 (Cell Signaling) for 1 h at room temperature. After washing in TBST (1X), membranes were revealed using Clarity Western ECL Substrate (Bio Rad) or Maxi Clarity Western ECL Substrate (BioRad) in a LAS-4000 Fujifilm system. Analysis of immunoblotting images was performed manually using Fiji software.

### Immunoprecipitation

For immunoprecipitation (IP) we used ORF1p (Millipore, MABC 1152) and IgG mouse (Thermofisher) antibodies. The antibodies were coupled to magnetic beads using the Dynabeads® Antibody Coupling Kit (Invitogen) according to the manufacturer’s recommendations. We used 8 µg of antibody for 1 mg of beads. The appropriate volume of buffer was added to the coupled beads to achieve a final concentration of 10 mg/ml.

LUHMES cells were plated at a density of 20×10^6^ cells in 2 T75 flasks and grown in differentiation medium for 5 days. One flask was treated with 50 µM arsenite for 1 h. Control and treated cells were washed with 1X PBS and harvested using 1 ml of lysis buffer (10 mM Tris HCl pH 8, 150 mM NaCl, NP40 0.5% v/v, protease inhibitor 10 µl/ml) per T75 flask. Samples were sonicated for 15 min at 4°C, then centrifuged at 1200 rpm for 15 min at 4°C. The supernatants obtained were transferred to a new microcentrifuge tube and then separated into 3 tubes. One tube containing 50 µl of supernatant (1/10th of the test samples) was stored at 4°C until the samples were analyzed. The other two tubes containing 445 µl of the remaining supernatants were used for IP. Mouse IgG beads (75 µl) were added to one tube and ORF1p antibody beads (75 µl) to the other tube. Each of these two tubes was then diluted to 1.5 ml with buffer (10 mM Tris HCl pH8, 150 mM NaCl, 10 µl/ml protease inhibitor) to dilute the NP40. The samples were then incubated overnight on a wheel at 4°C. Samples were then washed 3 times with 1 ml buffer (10 mM Tris HCl pH 8, 150 mM NaCl, 10 µl/ml protease inhibitor) using a magnet and then resuspended in 50 µl of the same buffer. The samples were boiled in Laemmli buffer (95°C, 10 min) and 20 µl of each sample were deposited on a 4-12% Nupage gel (Invitrogen). Migration was carried out in MES SDS running buffer (Invitrogen) for 45 min at 200 V. The gel was then transferred onto a PVDF membrane activated with methanol (Immobilon, Sigma) in glycine buffer (25 mM Tris, 200 mM glycine pH 8,3) for 1 h at 400 mA. The membranes were revealed using the primary antibodies anti-ORF1p rabbit 1:500 (Abcam, ab245249), anti-Lamin B1 rabbit 1:1000 (Abcam, ab16048), the secondary anti-rabbit HRP 1:2000 (Ozyme) with the Maxi Clarity Western ECL Substrate kit (BioRad). Analysis of immunoblotting images was performed manually using Fiji software.

### RT-qPCR

Total RNA from 5×10^5^ lysed cells was extracted using RNeasy Plus Micro Kit (Qiagen) according to the manufacturer’s instructions. The removal of contaminating genomic DNA (gDNA) was assessed by using the ezDNase kit (Invitrogen^TM^). After gDNA elimination, RNA (300 ng) was reverse transcribed using the All-In-One 5X RT MasterMix kit (Abcam) for first-strand cDNA synthesis ready for immediate real-time PCR analysis.

Quantitative PCR reactions were carried out in duplicates with SsoAdvanced Universal SYBR® Green Supermix (Biorad) on a CFX Opus 384 (Bio-Rad,) with the following program: enzyme activation at 95°C 2 min, primer denaturation at 95°C, annealing at 57°C and primer extension at 72°C for 42 cycles. The following primers were used: HPRT (F: CAGCCCTGGCGTCGTGATTAGT; R: CCAGCAGGTCAGCAAAGA AT); TBP (F: CAGCATCACTGTTTCTTGGCGT; R: AGATAGGGATTCCGGGAGTCAT) ; LINE-1 5’UTR (F: GTACCGGGTTCATCTCACTAGG R: TGTGGGATATAGTCTCGTG GTG), ORF1 (F: AGGAAATACAGAGAACGCCACAA) R: GCTGGATATGAAATTCTGGGTTGA), LINE-1 ORF2 (F: AAATGGTGCTGGGAAAACTG ; R: GCCATTGCTTTTGG TGTTTT), 3’UTR (F : GGGAATTGAACAATGAGATCAC; R: TATACATGTGCCATGC TGGTG). Data were analyzed using the ddCt method and values normalized to hypoxanthine-guanine phosphoribosyl transferase (HPRT) or to glyceraldehyde-3-phosphate dehydrogenase (TBP) TATA-binding protein.

### Proximity Ligation Assay (PLA)

The detection of the interactions between ORF1p and Lamin B1, NUP153 and KPNB1 were analyzed using proximity ligation Assay (PLA) method using Duolink® In Situ Red Starter Kit Mouse/Rabbit (Sigma, DUO92101). Cells were treated according to the manufacturer’s instructions. After PLA, cells are mounted with a Fluoromount (Invitrogen™, 15586276). Fluorescence images were acquired using a Spinning Disk W1 Yokogawa imaging system. The quantification was again conducted with the custom-written plugin (https://github.com/orion-cirb/Nuclei_ORF1P) described previously. In short, nuclei and cytoplasms are segmented in 3D using Cellpose. Within each cell, PLA foci are detected using Stardist 2D on each slice of the cell mask and then reconstructed in 3D. The foci parameters are ultimately computed in each cellular compartment (cytoplasm, nucleus, outer and inner nuclear membranes, and nucleoplasm).

### Immunofluorescence

Immunostaining experiments on cells were assessed at 1 h after arsenite treatment. Stressed and control differentiated LUHMES cells cultured in 24-well plates (150 000 cells/well) were fixed with 4% paraformaldehyde (PFA) in PBS for 20 min. Immunofluorescence were performed overnight at 4°C after a 20 min permeabilization in 0,2% Triton X100 in PBS and incubation in blocking buffer (10 % Normal goat serum (NGS), 0,1% Triton X-100 in PBS) for 1 h with the following antibodies: ORF1p 1:500 (Millipore, MABC1152 clone 4H1), TH 1:500 (Millipore, AB9702), Tuj1 1:1000 (Abcam, ab18207), Lamin B1 1:1000 (Abcam, ab16048), KPNB1 1:500 (GTX, 133733), NUP153 1:1000 (Novus biological, NB100-93329), H4K20me3 1:1000 (Abcam, ab9053), HP1 1:1000 (Cell Signaling, #2623S), TDP-43 1:600 (Proteintech, 107-82-2AP), HA 1:1000 (Roche, 11867423001), RanGAP1 1:500 (Santa Cruz, sc-28322), γH2AX 1:1000 (Abcam, ab2893), ssDNA 1:500 (ENZO, (F7-26) ALX 804-192) diluted in the blocking buffer, overnight at 4°C. For ssDNA immunostaining, cells were fixed with PFA, followed by 24 h methanol incubation at −20°C, then incubated at 37°C for 4 h with RNaseA. Fluorescence secondary antibodies: Alexa Fluor 555, Alexa Fluor 488, Alexa Fluor 647, 1:2000 (ThermoFisher Scientific) were added for 1 h at room temperature. Nuclei were stained with Hoechst 1:2000 (Invitrogen, H3570). We used Fluoromount (Invitrogen™, 15586276). Cells were imaged by Spinning Disk W1 Yokogawa and STED (stimulated-emission-depletion) Abberior Facility line imaging systems.

For immunostaining of post-mortem human brain tissues, sections were fixed with 4% paraformaldehyde (PFA) for 10 min. Sections were permeabilized in 0,5% Triton X-100 in PBS for 20 min and incubated at 100°C for 20 min in citrate buffer (10 mM citrate buffer pH 6.0, 0.05% Tween 20). After blocking for 1h (10 % Normal goat serum (NGS), 0,1% Triton X-100, in PBS), tissues were incubated overnight at 4°C with the following primary antibodies: ORF1p 1:500 (Millipore, MABC1152 clone 4H1), Lamin A/C 1:800 (Abcam, ab108595), NeuN 1:1000 (GENTEX, GTX00837) diluted in blocking buffer. The following secondary antibodies were added for 1 h at room temperature: Alexa Fluor 555, Alexa Fluor 488, Alexa Fluor 647 1:2000 (ThermoFisher Scientific). Nuclei were stained with Hoechst 1:2000 (Invitrogen, H3570). Sections were treated with TrueBlack lipofuscin autofluorescence quencher (OZYME, BTM23014) diluted in PBS for 10 min in the absence of light. The slides were rinsed 3 × 10 min in PBS to remove excess quencher and mounted with Fluoromount (Invitrogen™, 15586276). Sections were imaged by Spinning Disk W1 Yokogawa imaging system.

### Quantification and image analysis

The quantification of ORF1p fluorescence was performed with a custom-written plugin (https://github.com/orion-cirb/Nuclei_ORF1P) developed for the Fiji software ^24^, using Bio-Formats ^25^ and 3D Image Suite ^26^ libraries. Hoechst channel is downscaled by a factor 2, before detecting nuclei with the 2D-stitched version of Cellpose ^27^ (model = ‘cyto2’, diameter = 100, flow threshold = 0.4, cell probability threshold = 0.0, stitching threshold = 0.5). Segmented image is rescaled to original size and obtained 3D nuclei are filtered by volume to avoid false positive detections. Cells in ORF1p channel are detected in the same manner as nuclei. Then, each cell is associated with a nucleus having at least half of its volume in contact with it. Cells without any associated nucleus are filtered out. For each cell, ORF1p signal is analyzed in the following cellular compartments: cytoplasm, nucleus, outer and inner nuclear membranes and nucleoplasm (Fig. 2C, E). Compartmentalization is obtained using combinations of morphological operations (dilation, erosion and subtraction) of the cell and nucleus masks.

**Figure 2.**
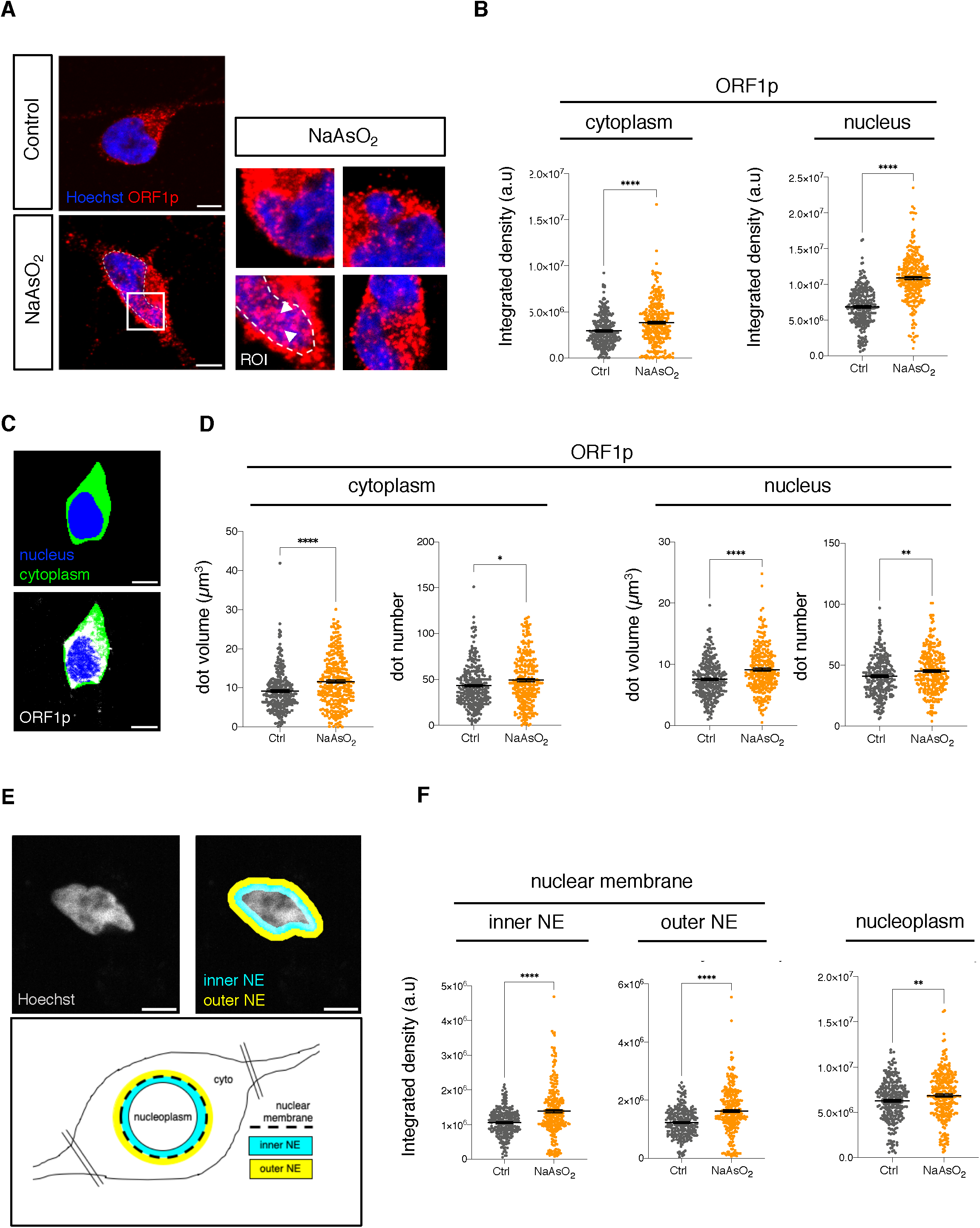
ORF1p increases, changes its conformational pattern and translocates to the nucleus following oxidative stress. (A) Immunostaining for LINE-1 ORF1p of differentiated LUHMES cells treated with arsenite. (B) Quantification of ORF1p intensity both in the cytoplasm and the nucleus in the presence and the absence of stress. n > 100 neurons were quantified per condition, a representative experiment (3 wells per condition) of 8 independent experiments, mean ± SEM, two-tailed t test (C) Scheme of the script (for details see Materials and Methods) for ORF1p quantification in the cytoplasmic, nuclear (nucleoplasm = nucleus – inner NE) and inner and outer NE compartments. (D) Quantification of the number and volume of ORF1p positive dots shown in ROI (region of interest) (A) in the cytoplasm and the nucleus using the same script, n > 100 neurons were quantified per condition, a representative experiment (3 wells per condition) of 8 independent experiments, mean ± SEM, two-tailed t test. (F) Quantification of ORF1p signal at the NE by an in-house script as schematized in (E) (for details see Materiel and Methods) shows increased ORF1p intensity at the inner NE, outer NE and in the nucleoplasm, n > 100 neurons were quantified per condition, a representative experiment (3 wells per condition) of 5 independent experiments, mean ± SEM, two-tailed t test (*p < 0.5, **p < 0.01, ***p < 0.001, ****p < 0.0001. Scale bar, 5 µm.

In each compartment, the integrated intensity of ORF1p is measured and normalized to ensure comparability across images. Normalization is done by subtracting the image background noise, estimated as the median intensity value of the minimum intensity Z-projection of the ORF1p channel. ORF1p foci are detected with Stardist 2D ^28^, applying a model trained on a dataset of 1950 images of foci. The following settings are used: percentile normalization = [0.2 - 99.8], probability threshold = 0.2, overlap threshold = 0.25. ORF1p foci are then labeled in 3D and filtered by volume. The number of foci, their volume and their normalized integrated intensity are computed in each cellular compartment. Results are provided as an Excel file.

To quantify nuclear deformation, another custom plugin (https://github.com/orion-cirb/HA_ORF1P_Prot/) was developed for the Fiji software, that uses the same libraries listed above, CLIJ ^29^ and Find Focused Slices (https://sites.google.com/site/qingzongtseng/find-focus?authuser=0). In brief, best focussed slice in the stack is selected. Nuclei are detected on this slice with Cellpose 2D (model = ‘cyto2’, diameter = 120, flow threshold = 0.4, cell probability threshold = 0.0), filtered by area and their circularity is computed. For each nucleus, ORF1p integrated intensity is analyzed in the following compartments: nucleus, outer and inner nuclear membranes and nucleoplasm. Compartments are obtained with the same combinations of morphological operations than previously described.

In case a HA-ORF1p channel is provided, cells are detected on the best focused slice with Cellpose 2D (model = ‘cyto2’, diameter = 200, flow threshold = 0.4, cell probability threshold = 0.0) and filtered by area. Nuclei having at least one pixel in contact with a HA-ORF1p cell are considered as HA-ORF1p+, other nuclei as HA-ORF1p-. The quantification of the intensity of other markers than ORF1p was also performed with this plugin.

The analysis of human sample images was conducted using a third custom-written plugin (https://github.com/orion-cirb/Hoechst_ORF1p_pTAU/) developed for the Fiji software, incorporating Bio-Formats and 3D Image Suite libraries. Nuclei detection in the Hoechst channel is performed using the 2D-stitched version of Cellpose (model = ‘cyto’, diameter = 100, flow threshold = 0.4, cell probability threshold = 0.0, stitching thresh-old = 0.5). The resulting 3D nuclei are filtered based on volume, and measurements of nuclei volume, circularity, as well as ORF1p intensity mean and standard deviation are conducted. Circularity is assessed on the middle plane of the nucleus, and intensity measurements are normalized by subtracting the image background noise, as described earlier.

If a NeuN channel is provided, NeuN-positive cells detection follows the same process as nuclei detection, with the exception that the model used was trained through the human-in-the-loop procedure implemented in the Cellpose GUI ^30^. Specifically, the model was initialized from the ‘cyto2’ pretrained Cellpose model and trained on 20 images (approximately 50 cells annotated). During the inference phase, the following parameters are employed: diameter = 140, flow threshold = 0.4, cell probability threshold = 0.0, stitching threshold = 0.5. Subsequently, each cell is linked to a nucleus with at least a quarter of its volume in contact with it. Nuclei are thus classified as either NeuN+ or NeuN-.

### Statistical Analysis

Data were obtained from independent experiments. The number of cells analyzed in each experiment is described in corresponding figure legends. Data analysis was performed using GraphPad Prism 8. Shapiro Wilk normality tests were performed prior to the statistical test to inform whether a parametric or non-parametric test should be used. Statistical tests used are indicated in the figure legends. The significant threshold was defined as p<0.5 and depicted as follows on all graphs: *p < 0.5, **p < 0.01, ***p < 0.001, ****p < 0.0001.

## Results

### Stress-induced activation of LINE-1 in human post-mitotic dopaminergic neurons

LUHMES cells represent a robust model to investigate the molecular and cellular mechanisms leading to neurodegeneration ^31–33^. LUHMES cells were differentiated into post-mitotic neurons according to the experimental paradigm ^21–23^ displayed in Figure 1A (and Suppl. Fig. 1A). These cells acquire a mature dopaminergic neuronal phenotype as revealed by immunostaining for the neuronal marker TuJ1 (β-III tubulin) and the dopaminergic marker TH (Tyrosine hydroxylase) at day 5 of differentiation as shown in Figure 1B (and quantified in Suppl. Fig. 1B). The dopaminergic phenotype of these cells was further confirmed by analyzing TH protein expression by Western blot at different days of differentiation (Suppl. Fig. 1C). Surprisingly, but as previously observed in mouse dopaminergic neurons *in vivo* ^5^, full-length LINE-1 elements are expressed at steady-state in cultured human dopaminergic neurons as revealed by immunostaining for the LINE-1 ORF1p protein (Fig. 1B). Human ORF1p, as mouse Orf1p ^5^, shows a predominant cytoplasmic expression pattern but is also observed nuclear. Oxidative stress is a common feature of aging ^34–36^, cancer ^37^ and NDs ^38–41^ and has been associated with LINE-1 activation ^42–44^. To generate a human neuronal model of endogenous LINE-1 activation, we induced oxidative stress in differentiated dopaminergic neurons with low-dose arsenite (NaAsO_2_ ; 50 µM) for 1 hour at day 5 of differentiation as described in Figure 1A. This treatment did not affect the neuronal and dopaminergic phenotypes as shown by TuJ1 and TH immunofluorescence quantification (Suppl. Fig. 1D, E) and triggered no apparent cell death. Oxidative stress increased the expression of LINE-1 in human neurons as revealed by Western blot and immunofluorescence analysis of LINE-1 ORF1p protein (Fig. 1C, D). LINE-1 RNA quantification by RT-qPCR using several primers targeting different regions of the LINE-1 sequence also showed an up-regulation of LINE-1 transcripts (Suppl. Fig. 1F). In order to test LINE-1 encoded protein activity, we analyzed two known consequences of ORF2p activity (for which no effective antibodies are available). Increased LINE-1 ORF2p has been previously reported to induce DNA damage due to the endonuclease activity ^3–5^. Upon oxidative stress, differentiated human dopaminergic neurons displayed increased DNA damage as quantified by an increase in γH2AX (phosphorylated histone H2AX)-positive foci, which decreased in the presence of stavudine, an inhibitor of the reverse transcriptase activity of LINE-1 ORF2p (Fig. 1E). The reverse transcriptase activity of LINE-1 ORF2p has also been reported to generate the accumulation of single-stranded DNA (ssDNA) in the cytosol leading to inflammation ^6,45^. We observed an increase of ssDNA foci volume and intensity in the cytosol of stressed neurons (Suppl. Fig. 2A). Full-length LINE-1 and its encoded proteins are thus expressed at steady-state in human dopaminergic neurons, are inducible by oxidative stress and functionally active. This *in vitro* model therefore represents a valuable tool to investigate cellular effects of LINE-1 activation mediated by the encoded proteins.

### ORF1p localizes to the nuclear envelope and translocates into the nucleus upon oxidative stress

We next investigated the precise cellular localization and expression levels of ORF1p upon stress using confocal imaging (Fig. 2A). ORF1p was predominantly cytoplasmic both in steady-state as well as in stress conditions and showed both, diffuse and dot-like patterns. Surprisingly and against the current assumptions ^46^, we observed the presence of ORF1p in the nuclear compartment and an accumulation of ORF1p at the nuclear periphery both at the outer and the inner NE as distinguishable by Hoechst DNA staining (Fig. 2A, B). We further quantified ORF1p intensities using an automated deep-learning based quantification script developed in-house as schematized (Fig. 2C). This script allows for the quantification of immunofluorescence intensities, dot numbers and volumes in pre-defined cellular compartments like the nucleus and the cytosol. The increase in staining intensities in both compartments was accompanied by a rearrangement of staining patterns as revealed by an increase in ORF1p-positive dot numbers and volumes, both in the cytoplasm and the nucleus (Fig. 2D), indicating a possible condensation of LINE-1 ORF1p in stress conditions. As ORF1p appeared to accumulate along the NE, the script was further applied to analyze ORF1p intensity in more detailed compartments including the outer NE, the inner NE and the nucleoplasm (not including the inner NE) (Fig. 2E). The quantification showed a significant increase of ORF1p in both the inner and outer NE and in the nucleoplasm (Fig. 2F).

In summary, ORF1p immunostainings using an unbiased image quantification method revealed increased LINE-1 ORF1p levels in the cytoplasm, in the nucleus, and in the inner and outer NE, indicating a general increase of ORF1p, the accumulation of ORF1p at the NE, a rearrangement of ORF1p in a dot-like manner particularly in the cytoplasm and a significant nuclear translocation upon stress.

### ORF1p targets nuclear envelope components upon oxidative stress

To confirm ORF1p localization to the inner NE, we performed co-immunostainings for ORF1p and Lamin B1, a nuclear lamina protein which decorates the inner NE and plays a key role in the maintenance of nuclear architecture. These experiments revealed the colocalization of ORF1p and Lamin B1 at the inner NE which increased upon stress. This colocalization is particularly visible at the level of lamina invaginations (Fig. 3A). Co-immunoprecipitation experiments confirmed the interaction between ORF1p and Lamin B1 under stress conditions (Fig. 3B). Direct binding of ORF1p to Lamin B1 was further examined by proximity ligation assay (PLA). The results obtained indicate a direct interaction between ORF1p and Lamin B1 at the nuclear periphery upon stress (Fig. 3C) in line with the colocalization signals observed by co-immunostainings (Fig. 1A). However, direct interaction of ORF1p and Lamin B1 is also detected in the nucleoplasm of neurons and PLA signal quantification confirmed an increase of the number of dots (= direct interaction events) at the inner NE but also in the nucleoplasmic compartment (Fig. 3D). This suggests a possible scenario in which the stress-induced nuclear translocation of ORF1p and direct ORF1p binding to Lamin B1 induces Lamin B1 mislocalization from the inner NE to the nucleoplasm.

**Figure 3.**
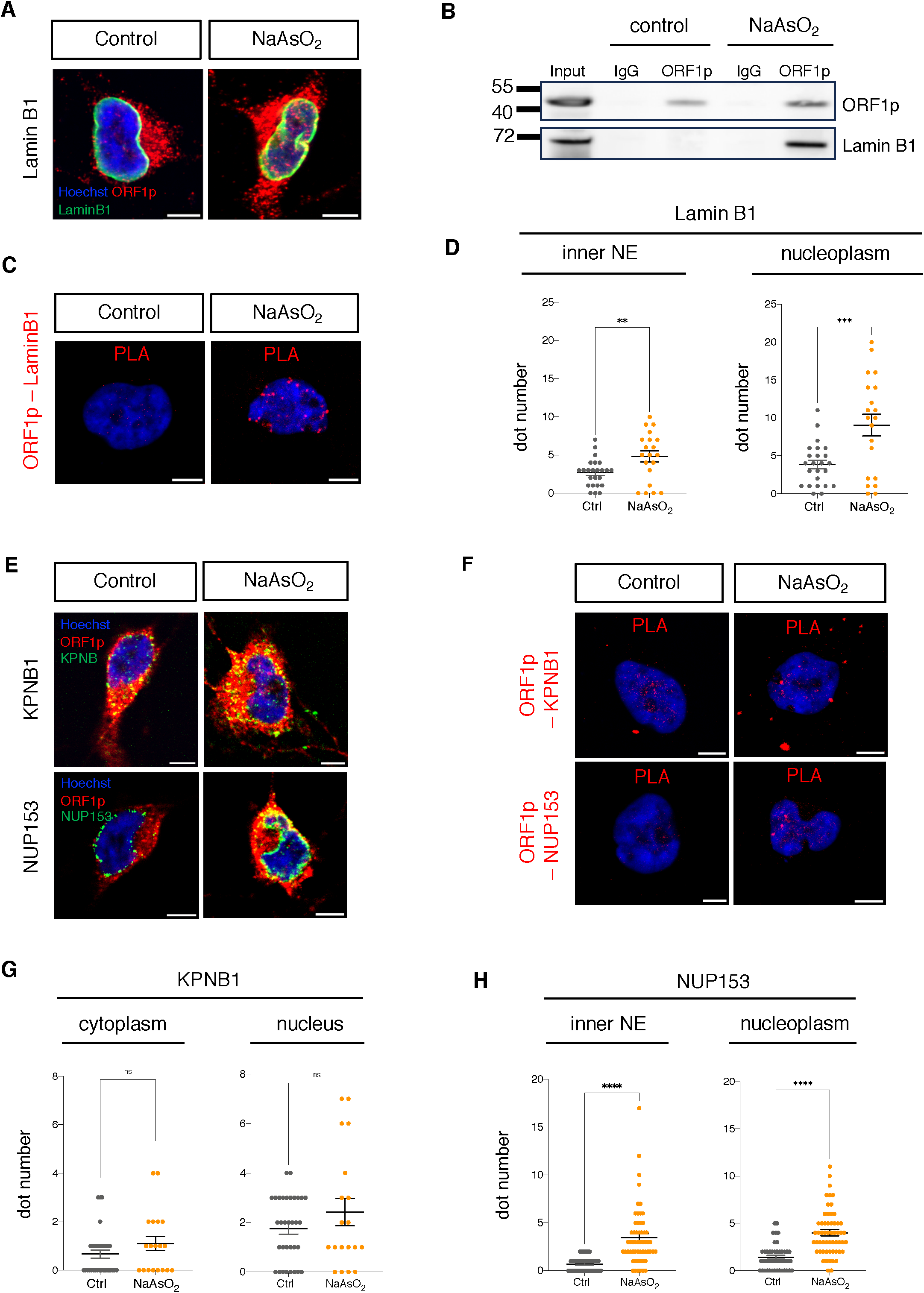
ORF1p association with nuclear envelope components and nuclear transport proteins increases upon oxidative stress. (A) Co-immunostaining of ORF1p and Lamin B1 reveals their colocalization (yellow). (B) Validation of ORF1p interaction with Lamin B1 by co-immunoprecipitation and Western-blotting. (C) Direct interaction of ORF1p with Lamin B1 was revealed by PLA and quantified in (D), n > 50 neurons were quantified per condition, (3 wells per condition). (E) Co-immunostaining of ORF1p and KPNB1 or NUP153. (F) Validation of a direct interaction between ORF1p and KPNB1 or NUP153 by PLA. The quantification in separate cellular compartments is shown in (G, H), n > 50 neurons were quantified per condition, (3 wells per condition), mean ± SEM, two-tailed t test (*p < 0.5, **p < 0.01, ***p < 0.001, ****p < 0.0001). Scale bar, 5 µm.

Previous studies have shown that ORF1p interacts with importin beta family members (*i.e*, KPNB1) ^47,48^, which could mediate its nuclear translocation using nuclear pore complexes (NPCs). To confirm that nuclear translocation of ORF1p in neurons upon oxidative stress also mediated by this nuclear import pathway, we performed co-immunostaining of ORF1p with KPNB1 and NUP153 (a nucleoporin of the nuclear basket), a core NPC protein reported to interact with KPNB1 ^49,50^. The results shown in Figure 3E indicate the colocalization of ORF1p with KPNB1 and NUP153 at the NE indicating that nuclear translocation of ORF1p in human neurons upon oxidative stress might indeed be mediated by these proteins. In addition, several ORF1p-positive cytoplasmic foci colocalized with KPNB1 (Fig. 3E) which might jeopardize normal KPNB1 function. Direct interaction of ORF1p with KPNB1 and NUP153 was further confirmed by PLA (Fig. 3F). PLA signal of ORF1p and KPNB1 interaction is not affected by oxidative stress (Fig. 3G), whereas the direct interaction between ORF1p and NUP153 is increased in stress condition as indicated by increased number of PLA foci at the inner NE and the nucleoplasm (Fig. 3H). Altogether, these results reveal that ORF1p directly interacts with components of the inner NE and the nucleocytoplasmic transport (NCT) machinery and might interfere with their physiological functions in nuclear architecture and NCT function. This let us to investigate whether ORF1p localization to the NE and direct protein-protein interaction with Lamin B1, KPNB1 and NUP153 might affect NE morphology and/or NCT efficiency.

### Nuclear ORF1p accumulation correlates with nuclear envelope alterations

Lamin B1 staining of the inner nuclear lamina of human dopaminergic neurons revealed various NE dysmorphologies including nuclear invaginations and blebbing which appeared upon oxidative stress (exemplified in Fig. 4A). It is noteworthy that nearly 70% of stressed cells presented such deformations (Fig. 4B). Co-immunostaining for ORF1p and Lamin B1 revealed the accumulation of ORF1p at the level of NE invaginations and blebs, which is clearly observable in STED acquired 3D images (Fig. 4A). Nuclear dysmorphology was accompanied by increased Lamin B1 levels at the inner NE but even more so in the nucleoplasm of stressed cells (Suppl. Fig. 2B), again suggesting a delocalization of Lamin B1 possibly as a result of direct protein interaction with ORF1p. The degree of NE distortion (expressed as a circularity index) was analyzed using an automated script (detailed in the Materiel and Methods section) which measures nuclear circularity using Lamin B1 staining as a mask (Fig. 1D). This analysis showed a significant decrease of nuclear circularity in stressed cells (Fig. 4C). Importantly, the degree of nuclear distortion (the lower the index, the higher the distortion) was significantly correlated with the intensity of nuclear ORF1p staining (thus nuclear ORF1p quantity) both in the nucleoplasm (p<0,0001; Spearman r = −0,8230; 95% CI −0,8578-0,7807) and at the inner NE (p<0,0001; Spearman r = −0,9169; 95% CI −0,9339-0,8958, Fig. 4E). This correlation suggests that nuclear ORF1p translocation and accumulation in the nucleoplasm and at the inner NE might be involved, at least partly, in the occurrence of nuclear dysmorphologies observed in neurons upon oxidative stress.

**Figure 4.**
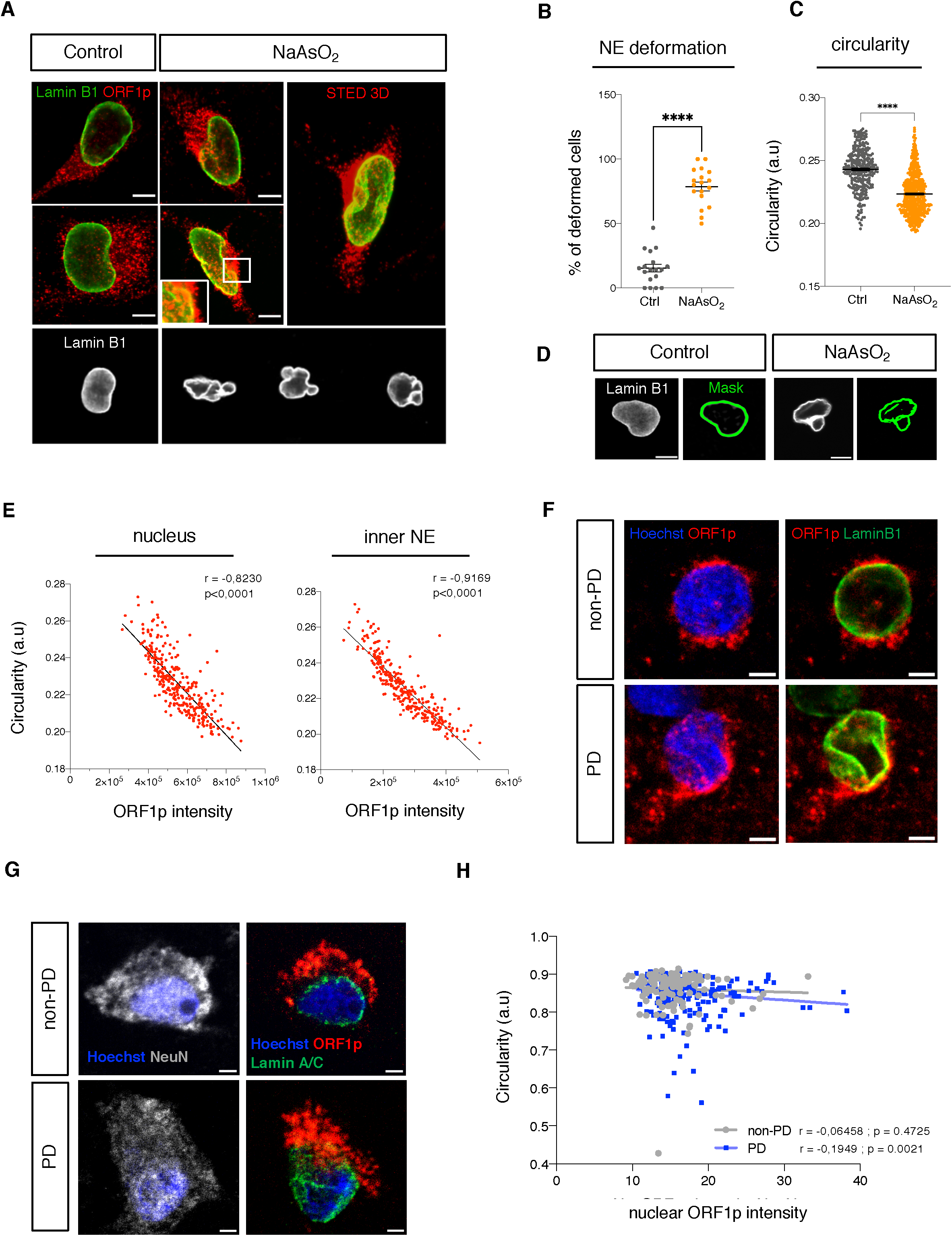
Stress induced association of ORF1p to the nuclear envelope leads to nuclear envelope dysmorphology. (A) NE dysmorphology in the arsenite stress model revealed by co-immunostaining of ORF1p and LaminB1 quantified in (B). The percentage of deformed cells was quantified in > 10 images per condition, a representative experiment (3 wells per condition) of 3 independent experiments is shown, mean ± SEM, two-tailed t test. (C) Quantification of nuclear dysmorphology by analyzing nuclear circularity using a dedicated script (for details see Materiel and Methods) based on Lamin B1 as a mask (D), shows decreased circularity under stress conditions. n > 100 neurons were quantified per condition, a representative experiment (3 wells per condition) of 5 independent experiments, mean ± SEM, two-tailed t test. (E) Correlation of ORF1p intensity per cell in the nucleus or at the inner NE with nuclear circularity. n > 100 neurons were quantified per condition, a representative experiment (3 wells per condition) of 3 independent experiments; Spearman correlation (*p < 0.5, **p < 0.01, ***p < 0.001, ****p < 0.0001). (F) Co-immunostaining of ORF1p and Lamin A/C in neurons of the cingulate gyrus of post-mortem PD and non-PD tissues. (G) Immunostaining of NeuN, Lamin A/C and ORF1p in the cingulate gyrus of post-mortem PD and non-PD tissues. (H) Correlation of nuclear ORF1p intensity and circularity per NeuN-positive cell in the cingulate gyrus of post-mortem tissues from PD patients (n = 5) and non-PD affected individuals (n = 4); Spearman correlation (*p < 0.5, **p < 0.01, ***p < 0.001, ****p < 0.0001).

Human dopaminergic neurons are particularly vulnerable to neurodegeneration observed in PD ^51^, express ORF1p more abundantly than surrounding neurons and their degeneration can be driven by LINE-1 expression in mice ^5^. In addition, NE alterations have been previously described in dentate gyrus brain tissues of sporadic and LRRK2-linked PD samples ^52^. In order to establish whether ORF1p is expressed in neurons in the human brain, and if so, whether nuclear ORF1p levels correlate with NE deformations, we turned to human post-mortem brain tissues of PD patients and non-PD affected controls some of whom had signs of early AD pathology (Braak 1-2; Table 1). We chose the cingulate gyrus, a region secondarily affected in PD, reasoning that dopaminergic neurons of the *substantia nigra pars compacta*, were most probably already lost at these advanced stages of the disease. We performed immunostainings using ORF1p, the pan-neuronal marker NeuN and either Hoechst to identify the nucleus or Lamin A/C to identify the NE and used an unbiased machine-learning assisted cell detection workflow which (a) identifies neurons based on the presence of NeuN, (b) delimits the nucleus based on Hoechst staining, (c) quantifies ORF1p within this delimitation and (d) measures the circularity of the nuclei. Immunostainings of ORF1p and Lamin A/C or Hoechst show a mostly cytoplasmic but also nuclear localization of ORF1p in neurons in samples from both, PD patients and non-PD controls (Fig. 4F, G). Importantly, analysis of a correlation of nuclear ORF1p intensity with nuclear circularity in each identified neuron identified a significantly negative correlation between the two parameters in PD samples (p=0.0021; Spearman r= −0.1949; 95% CI −0.32-0.07) but not in controls (p= 0.4725; Spearman r = −0.06458; 95% CI −0.24-0.12, Fig. 4H). As in human dopaminergic neurons in culture, this suggests a relationship in human post-mortem neurons in which the higher the concentration of ORF1p, the more pronounced the deformation of the nucleus.

**Table 1.** List of PD and non-PD post-mortem tissues used in this study.

| Internal ID | Diagnosis | PMI (hr) | Gender | Age at death | Pathological staging |
| --- | --- | --- | --- | --- | --- |
| 6629 | Parkinson Disease | 35 | M | 72 | Diffuse Lewy Body Disease |
| 8418 | Parkinson Disease | 36 | M | 82 | Diffuse Lewy Body Disease |
| 7826 | Parkinson Disease | nd | F | 62 | Diffuse Lewy Body Disease |
| 6532 | Parkinson Disease | 48 | F | 94 | Diffuse Lewy Body Disease |
| 3771 | Parkinson Disease | 20 | M | 83 | Diffuse Lewy Body Disease |
|  |  | 34.8 (+/-11.2) | 2F / 3M | 78.6 (+/-12.1) |  |
| 6203 | Non-PD | 23 | M | 78 | Alzheimer lesion Braak II |
| 8866 | Non-PD | 15 | M | 84 | - |
| 4078 | Non-PD | 10 | M | 73 | Alzheimer lesion Braak II |
| 5603 | Non-PD | 21 | F | 92 | Alzheimer lesion Braak I |
| 7197 | Non-PD | 10 | M | 85 | Alzheimer lesion Braak II |
|  |  | 15.8 (+/-6.1) | 3F/ 4M | 82.4 (+/-7.2) |  |

PMI, post-mortem interval.

In an attempt to establish causality between an increase in nuclear ORF1p and nuclear dysmorphologies, we used siRNA to knock-down LINE-1 expression. However, this approach did not significantly decrease nuclear ORF1p. We therefore turned to a different approach by blocking nuclear import of ORF1p using nuclear import inhibitors. As mentioned above, ORF1p nuclear import is mediated by a KPNB1/KPNA2-dependent process ^47,48^. KPNA2 is an importin alpha family member, which recognizes NLS of cargo proteins and serves as an adapter for KPNB1 for the import activity of the complex. KPNA2 has been shown to be involved in nuclear import of ORF1p in addition to KPNB1 ^47,48^. Using ivermectin, a KPNB1 inhibitor ^53,54^, and importazole, an inhibitor of KPNA2 ^55,56^, we obtained a significant decrease in nuclear ORF1p upon stress (Fig. 5A), which we did not observe with either drug alone (Suppl. Fig. 2C, D). ORF1p nuclear import in human neurons is thus dependent on both, KPNB1 and KPNA2. Importantly, the decrease in nuclear circularity upon stress was significantly rescued by blocking the import of ORF1p into the nucleus (Fig. 5A). Therefore, the correlation of an increase of nuclear ORF1p with a decrease in nuclear circularity as well as the restoration of nuclear deformations by blockade of ORF1p nuclear import in stress conditions strongly indicates that nuclear ORF1p causally participates in the loss of NE integrity. It is noteworthy that ORF2p reverse transcriptase inhibitor (stavudine) which prevented ORF2p induced DNA strand breaks (Fig. 1E) did not have any effect on nuclear circularity, or Lamin B1 delocalization, nor on ORF1p nuclear and cytoplasmic levels (Suppl. Fig. 3A-C). This indicates that the effect observed is not due to ORF2p activity.

**Figure 5.**
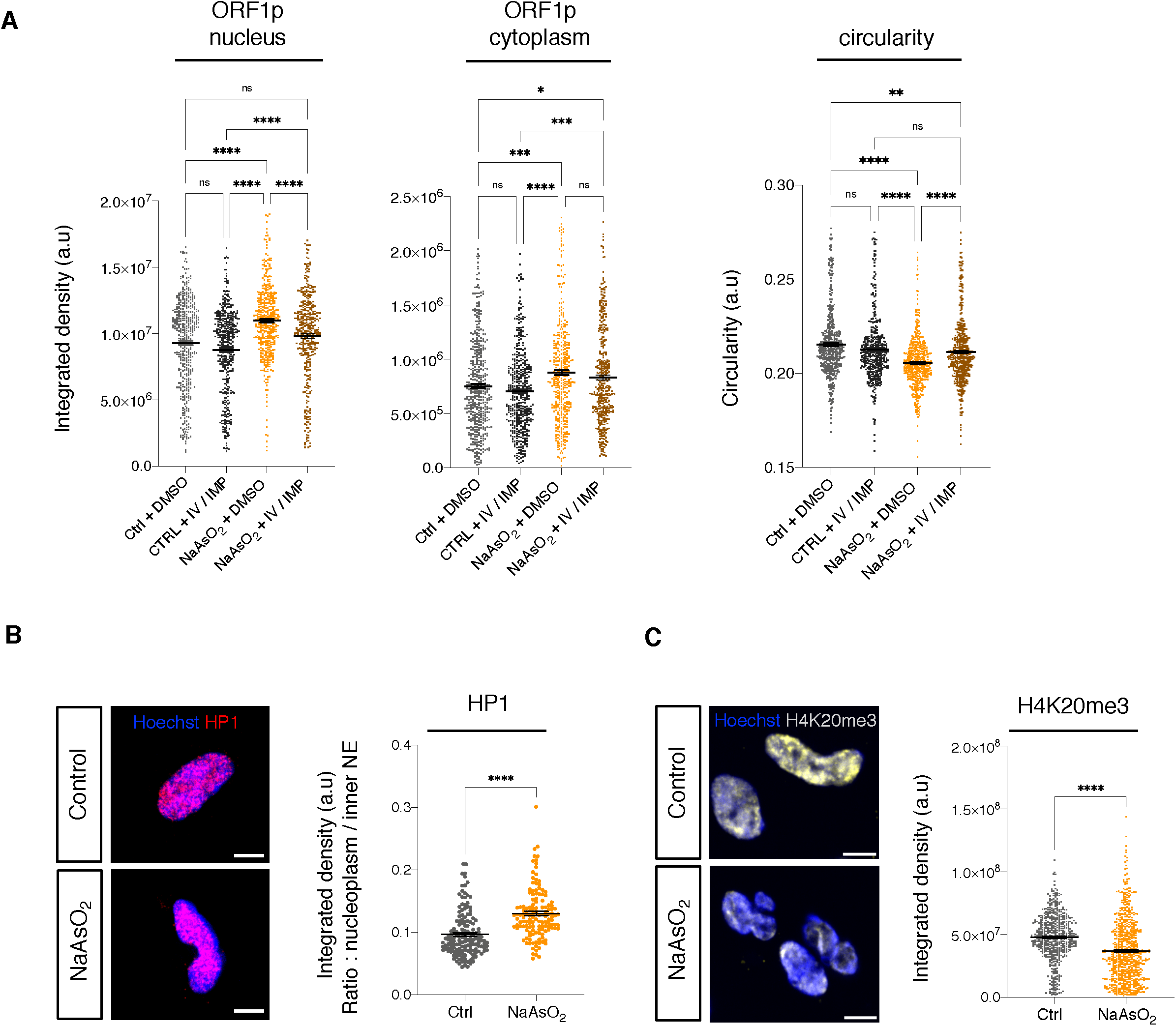
Restauration of nuclear circularity upon blockade of ORF1p nuclear import and analysis of chromatin repressive marks upon stress. (A) Quantification of ORF1p nuclear intensity and the nuclear circularity in the presence or the absence of import inhibitor drugs ivermectin (IV) and importazole (IMP) in stressed and control cells. n > 100 neurons were quantified per condition, a representative experiment (3 wells per condition) of 2 independent experiments, mean ± SEM, One-way ANOVA with Tukey’s multiple comparisons test. (B) Immunostaining of HP1 in stressed cells and the ratio of HP1 intensity in the nucleoplasm over the nuclear periphery (inner NE). n > 100 neurons were quantified per condition, a representative experiment (3 wells per condition) of 3 independent experiments, mean ± SEM, two-tailed t test. (H) Immunostaining of H4K20me3 repressive histone mark in stressed and control cells and its quantification in the nucleus. n > 100 neurons were quantified per condition, a representative experiment (3 wells per condition) of 3 independent experiments, mean ± SEM, two-tailed t test (*p < 0.5, **p < 0.01, ***p < 0.001, ****p < 0.0001). Scale bar, 5 µm.

Since nuclear lamina plays an important role in anchoring heterochromatin at the inner NE (= nuclear lamina), we examined whether the observed alteration of the nuclear lamina is accompanied by heterochromatin disorganization by immunostaining for the heterochromatin markers HP1 (heterochromatin protein-1) and the repressive histone mark H4K20me3 (histone H4 trimethylated on lysine 20) in control and stressed cells. HP1 was significantly depleted from the inner NE (Fig. 5B) and H4K20me3 levels decreased in stress conditions (Fig. 5C) suggesting possible decondensation or detachment of heterochromatin from the inner NE and favoring conditions allowing the activation of LINE-1 expression ^57^.

### Nuclear envelope targeting by ORF1p is accompanied by nucleocytoplasmic transport defects

In view of the interaction of ORF1p with KPNB1 and NUP153 reported above, we examined whether the accumulation of ORF1p at the NE could lead to NCT deficiencies. Immunostainings and subsequent quantification of intensity, distribution and localization of KPNB1 showed an accumulation of KPNB1 in the cytoplasm, as revealed by an increase in the intensity of KPNB1. We also observed a change in KPNB1 distribution, as revealed by an increase in the number of counted dots and their volume in the cytoplasm (Fig. 6A) indicating a possible alteration of its function. To examine whether these changes in KPNB1 upon stress were associated with NCT defects, we first analyzed the localization of a major nucleocytoplasmic transport protein, RanGAP1, by immunostaining. The GTPase activity of this protein is involved in maintaining the Ran gradient required for NCT and its delocalization is associated with NCT defects ^58^. RanGAP1 accumulated in the cytoplasm, shown by increased intensity, number and dot volume in the cytoplasm (Fig. 6B) which is indicative of an alteration of NCT. Disruption of NCT can lead to the abnormal localization of the RNA binding protein TDP-43 (TAR DNA-binding protein 43), a predominantly nuclear protein. Mutations in TDP-43 gene (*TARDBP*) are causal in familial forms of FTD (Frontotemporal dementia) and ALS and a change from its predominantly nuclear to cytoplasmic localization has been observed in several NDs ^59,60^. Current consensus suggests that TDP-43 misdistribution leads to a pathological gain-of-function in the cytoplasm and loss-of-function in the nucleus ^61,62^ contributing to cellular dysregulation culminating in neurodegeneration. We therefore performed immunostainings for TDP-43 which revealed the presence of TDP-43-positive foci in the cytoplasm under stress conditions which were absent in non-stressed controls. Quantification showed that the intensity as well as the number and volume of TDP-43 positive dots were increased in the cytoplasm in stressed neurons (Fig. 6C). NUP153 is a nucleoporin which is involved in NCT and its cytoplasmic mislocalization and/or intranuclear accumulation are associated with NCT defects in various neurodegenerative contexts ^58,63,64^. The quantification of NUP153 showed an abnormal intranuclear and cytoplasmic accumulation of NUP153 in stress conditions (Fig. 6D).

**Figure 6.**
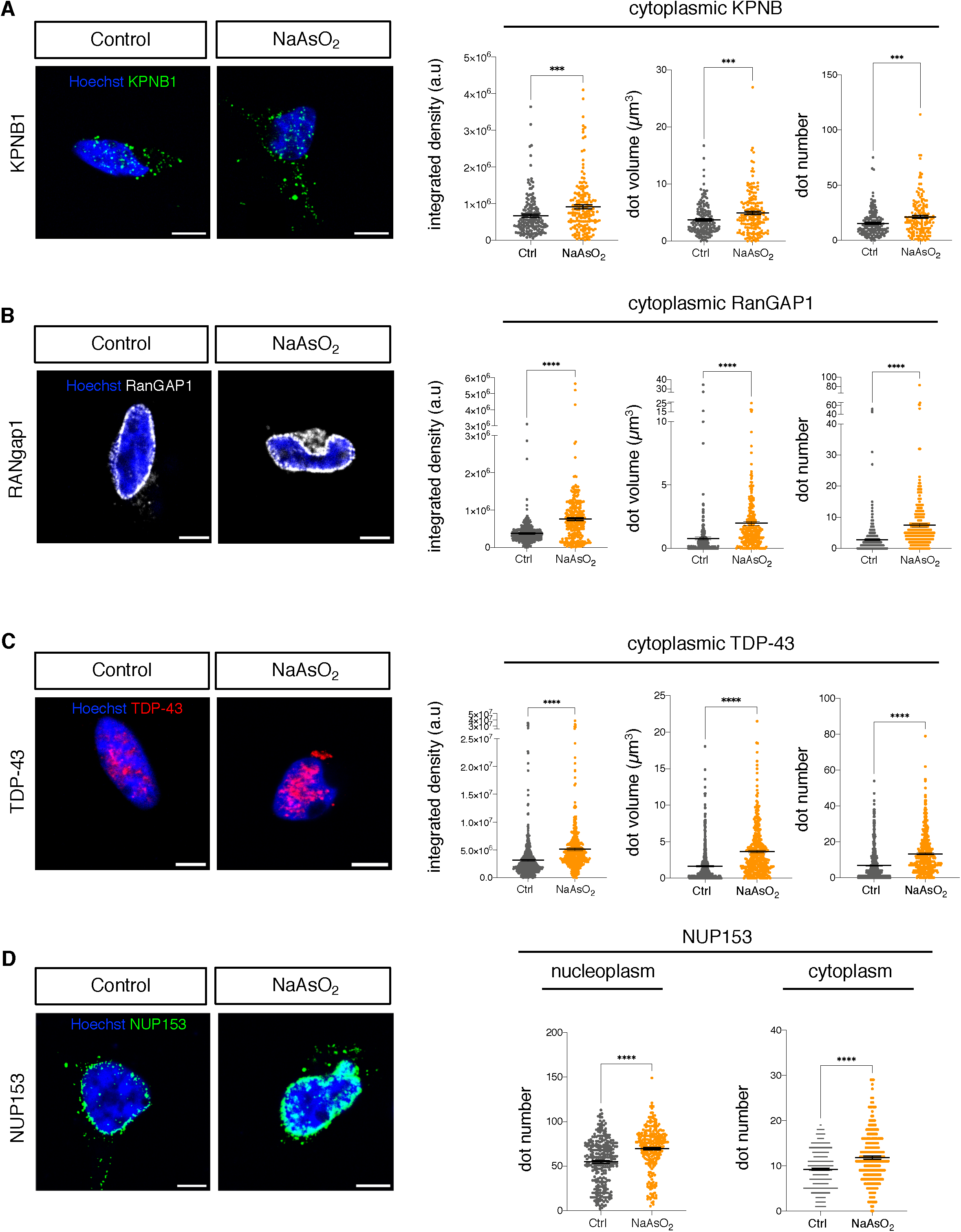
Stress induced association of ORF1p with the nuclear envelope is accompanied by nucleocytoplasmic transport defect. (A) Immunostaining of KPNB1 in oxidative stress conditions and quantification of the intensity levels, the number and the volume of dots in the cytoplasm. (B) Immunostaining of NPC protein RanGAP1 in oxidative stress conditions and quantification of the intensity levels, the number and the volume of dots in the cytoplasm. (C) Immunostaining of TDP-43 in oxidative stress conditions and quantification of the intensity levels, the number and the volume of dots in the cytoplasm. (D) Immunostaining of NUP153 in oxidative stress conditions and quantification of the intensity levels in the nucleoplasm and the cytoplasm. n > 100 neurons were quantified per condition, a representative experiment (3 wells per condition) of 3 independent experiments, mean ± SEM, two-tailed t test (*p < 0.5, **p < 0.01, ***p < 0.001, ****p < 0.0001). Scale bar, 5 µm.

### ORF1p overexpression leads to nuclear envelope dysmorphology and nucleocytoplasmic transport defects in the absence of oxidative stress

In order to evaluate whether NE dysmorphology induced by an increase in nuclear ORF1p could also be observed in the absence of arsenite-induced stress, we performed gain of function experiments by nucleofection of a plasmid expressing HA-tagged ORF1p but not ORF2p. Dopaminergic neurons were nucleofected at day 3 of differentiation with a plasmid encoding either HA-ORF1 or GFP control and analyzed 48 h later. Co-immunostainings for HA and Lamin B1 showed the presence of HA-ORF1 in the nucleus and at the inner NE, colocalizing with Lamin B1 (Fig. 7A). HA-ORF1 transfected cells displayed pronounced NE deformations as revealed by Lamin B1 staining (Fig. 7A), decreased circularity and increased nucleoplasmic Lamin B1 levels compared to GFP expressing cells (Fig. 7B). Interestingly, nuclear circularity was highly correlated with the nuclear intensity of HA-ORF1 in transfected cells (Fig. 7B), strongly suggesting a link between ORF1p nuclear localization and NE dysmorphology. Neurons transfected with HA-ORF1 also presented increased levels of RanGAP1 in the cytoplasm and TDP-43 in both the cytoplasm and nucleus compared to GFP expressing cells (Fig. 7C-F). These results indicate that overexpression of ORF1p, in the absence of stress, is sufficient to induce NCT defects and NE dysmorphology in human dopaminergic neurons and again establish a causal link between NE abnormalities and nuclear ORF1p content.

**Figure 7.**
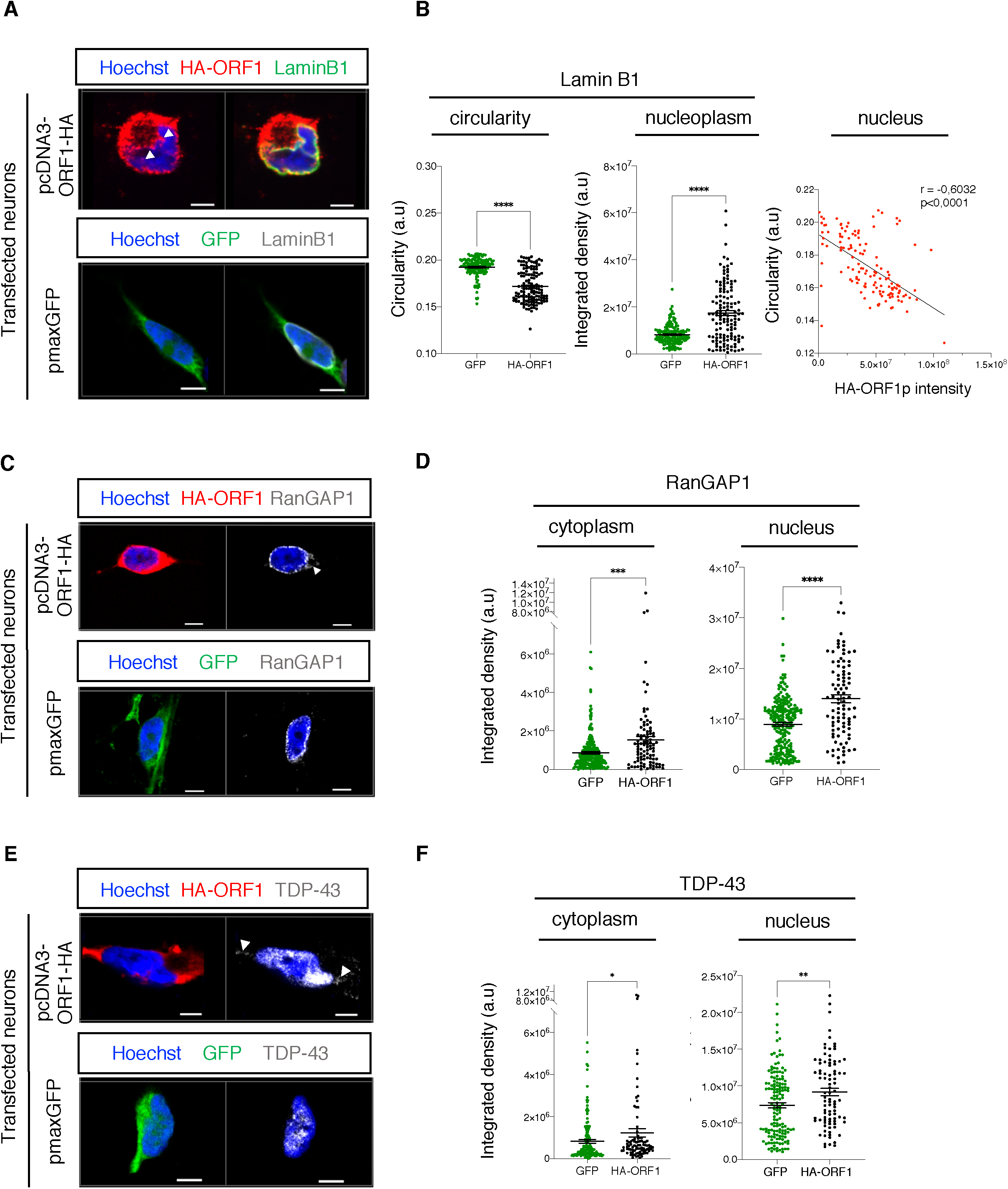
LINE-1 over-expression following transfection leads to nuclear envelope deformation and nucleocytoplasmic transport defects in the absence of stress. (A) Co-Immunostaining of HA and Lamin B1 in cells transfected with pcDNA3-ORF1-HA or with pmaxGFP as a control. (B) Quantification of the nuclear circularity index, Lamin B1 intensity in the nucleoplasm (left panel; mean ± SEM, two-tailed t test) and correlation of HA-ORF1p intensity per cell in the nucleus with nuclear circularity (right panel; Spearman correlation). n > 90 neurons were quantified per condition, a representative experiment (3 wells per condition) of 3 independent experiments; (*p < 0.5, **p < 0.01, ***p < 0.001, ****p < 0.0001). (C) Immunostaining of HA and RanGAP1 in cells transfected with pcDNA3-ORF1-HA or with pmaxGFP as a control. (D) Quantification of cytoplasmic RanGAP1 intensity in transfected cells with pcDNA3-ORF1-HA or pmaxGFP as a control. n > 90 neurons were quantified per condition (3 wells per condition), mean ± SEM, two-tailed t test. (E) Immunostaining of HA and TDP-43 in cells transfected with pcDNA3-ORF1-HA or with pmaxGFP as a control. (F) Quantification of cytoplasmic and nuclear TDP-43 signal in cells transfected with pcDNA3-ORF1-HA or pmaxGFP as a control. n > 100 neurons were quantified per condition (3 wells per condition), mean ± SEM, two-tailed t test (*p < 0.5, **p < 0.01, ***p < 0.001, ****p < 0.0001). Scale bar, 5 µm.

### Rescue of nuclear envelope dysmorphology under stress conditions by remodelin

NE morphology is disrupted during aging ^65^ and in premature aging syndromes caused by laminopathies ^66,67^ including progeria syndromes ^68^. In Hutchinson-Gilford progeria syndrome (HGPS), nuclear integrity was shown to be restored by the small molecule remodelin, an inhibitor of the acetylase-transferase protein NAT10 ^69^. But whether this molecule is efficient in restoring misshapen nuclear morphology in cell types other than fibroblasts or epithelial cells was not known. We therefore examined whether NE dysmorphology induced by ORF1p under oxidative stress in human neurons could be restored by remodelin. NE deformations (i.e invaginations) were clearly restored in cells pre-treated with remodelin for 24 h (Fig. 8A, B). Quantification showed that the decrease of nuclear circularity following arsenite treatment was restored by remodelin (Fig. 8B), which was paralleled by a reduction of nucleoplasmic Lamin B1 levels quantified in Figure 8B. In order to investigate whether remodelin treatment had an influence on ORF1p levels, we performed ORF1p immunostainings on cells treated or not with remodelin. Arsenite-induced nuclear ORF1p increase was completely normalized in the presence of remodelin, both at the inner NE and in the nucleoplasm (Fig. 8C) Interestingly, in these conditions cytoplasmic ORF1p levels remained unchanged at an expected higher level than non-stressed controls. Since remodelin has been reported to restore a “juvenile” chromatin state in Progeria models ^69,70^, we examined H4K20me3, known to repress LINE-1 elements ^57,71^. As shown in Figure 8A, H4K20me3 immunostaining was significantly decreased following stress and restored with remodelin treatment as confirmed by quantification of H4K20me3 intensity in the nucleus (Fig. 8D). This suggests that the decrease of nuclear ORF1p upon remodelin treatment in stressed cells might induce indirect blockade of nuclear ORF1p translocation. Alternatively, this decrease could also be due to a transcriptional inhibition of LINE-1 or a consequence of deposition of H4K20me3 at LINE-1 loci.

**Figure 8.**
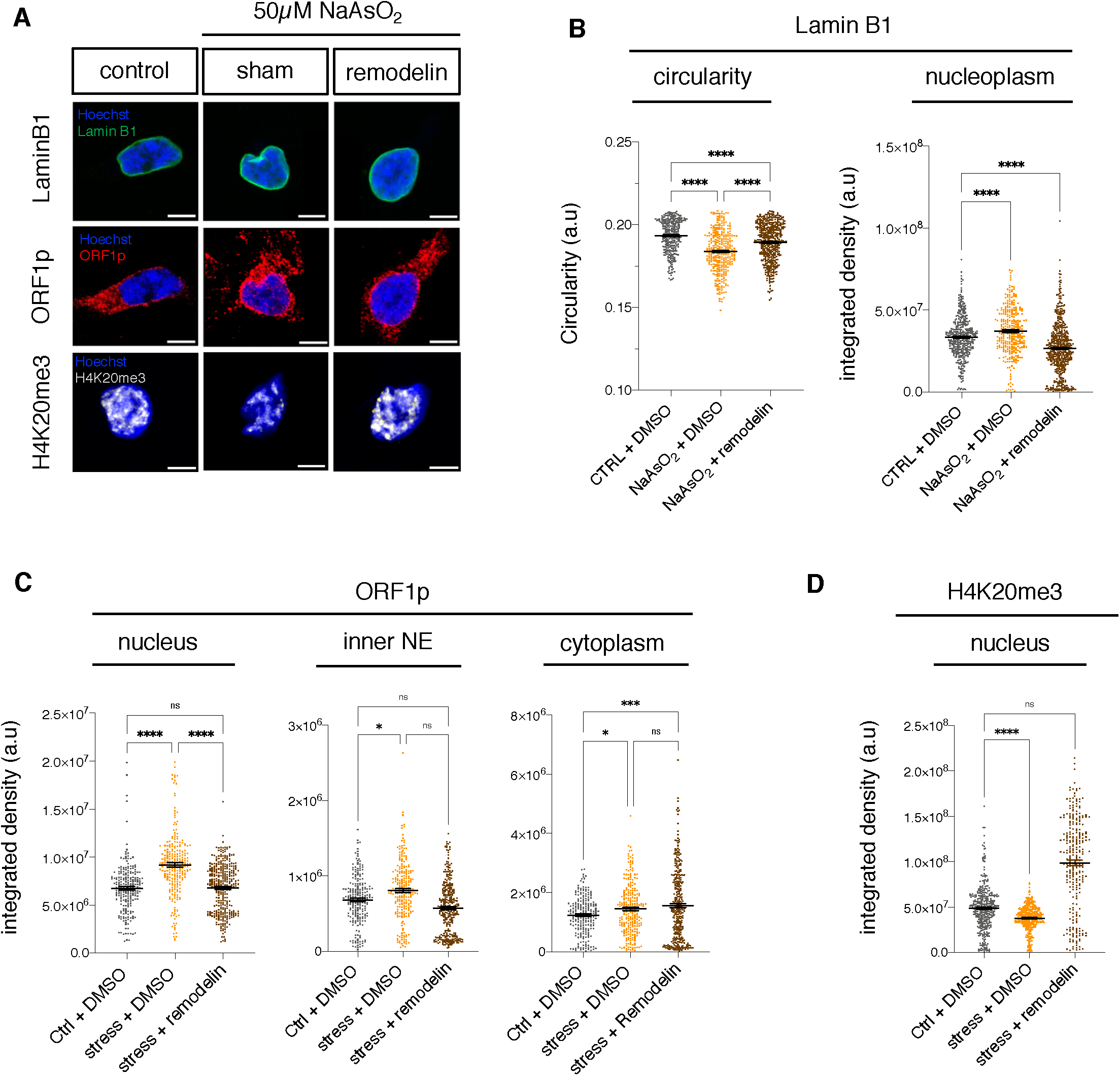
Restauration of nuclear alterations by the small molecule remodelin. (A) Immunostaining of ORF1p, Lamin B1 and H4K20me3 of neurons with or without remodelin pre-treatment 24 h before addition of arsenite or sham. (B) Quantification of nuclear circularity and Lamin B1 intensity in the nucleoplasm. (C) Quantification of ORF1p intensity in the nucleus, in the inner NE and in the cytoplasm. (D) Quantification of H4K20me3 intensity in the nucleus. n > 100 neurons were quantified per condition, a representative experiment (3 wells per condition) of 3 independent experiments, mean ± SEM, One-way ANOVA with Tukey’s multiple comparisons test (*p < 0.5, **p < 0.01, ***p < 0.001, ****p < 0.0001). Scale bar, 5 µm.

## Discussion

In recent years, TEs have emerged as possible new pathogenic players in NDs ^5,10,13,19,20,72–74^. However, the mechanisms through which LINE-1 might contribute to the pathogenesis of NDs still remain poorly understood potentially implicating cis-genetic or epigenetic, LINE-1 RNA or LINE-1 protein-mediated effects. Concerning the latter, the study of potential pathogenic effects of LINE-1 ORF2p has received most attention due to its enzymatic activities as an endonuclease and reverse transcriptase while ORF1p has widely been reduced to an RNA chaperone with cis-preference for its own RNA and a necessary component of the LINE-1 ribonucleoprotein particle (RNP) for retrotransposition. Previous work suggests that ORF2p is not only a source of genomic instability ^3–6,20^, but can also trigger the innate immune response through reverse transcription of RNA generating cytoplasmic ssDNA ^6,8,45^. While ORF2p reverse transcriptase activity can be targeted by inhibitors (nucleoside reverse transcriptase inhibitors, NRTIs, i.e., stavudine), the analysis of ORF2p protein expression and localization remains challenging due to a lack of efficient antibodies and/or its lesser abundance compared to ORF1p (1 ORF2p for 240 ORF1p molecules per LINE-1 RNP ^75^). In contrast, no inhibitor has been identified to target ORF1p activity, but effective antibodies are available to explore its expression and subcellular localization in pathological situations. However, a better understanding of the implication of ORF1p in the LINE-1 life cycle and beyond is important as accumulating data suggests that full-length LINE-1 are expressed, predominantly in epithelial cancers ^76^ but also at steady-state in mouse neurons ^5^, human neurons ^77^ and post-mortem brain tissues ^78^.

Using a human neuronal model of oxidative stress which displays increased LINE-1 activity as monitored by an increase in ORF1p expression and ORF2p-dependent genomic instability, we observed not only an increase of ORF1p in the cytoplasm, but also a striking translocation of ORF1p to the nucleus. Current consensus postulates that LINE-1 encoded proteins enter the nucleus only during cell cycle ^46^ which thus would exclude the presence of LINE-1 proteins in the nucleus of non-dividing cells including post-mitotic neurons. However, others have reported the presence of ORF1p in the nucleus of human cells in a neurodegenerative context ^79^ and in post-mortem human brain tissues ^78^. We also observed ORF1p staining in the nucleus of human dopaminergic neurons *in vitro* and in mouse dopaminergic neurons *in vivo* even at steady-state ^5^, and oxidative stress dramatically increases the nuclear translocation of ORF1p. The most probable mode of import for ORF1p is the importin-beta pathway. Indeed, ORF1p, as we show here in neurons and others have shown in HeLa cells ^47^ binds directly to the importin alpha and beta family members, KPNA2 and KPNB1, which are transport proteins carrying cargo proteins through the NPC. This importin-assisted mode of nuclear entry is widely used by several viral proteins ^80,81^, certain retrotransposons (i.e Ty1 ^82^, LINE-1 ORF1p itself ^48^) but also a pathological form of Tau which aggregates in neurodegeneration ^83^. The binding of ORF1p to KPNB1 does not increase under stress condition indicating that the stress-induced increase of ORF1p nuclear translocation probably requires other protein partners. It is noteworthy that ORF1p, in a stress-dependent manner, accumulates both at the outer NE and the inner NE which corresponds to the nuclear lamina and co-localizes with and binds directly to the NPC protein NUP153. This suggests a NUP153-dependent nuclear import of ORF1p under stress.

The small transcriptional increase in LINE-1 RNA following arsenite treatment suggests that the overall increase in ORF1p could be partially transcriptional and partially post-transcriptional, possibly by releasing LINE-1 RNA stored in P-bodies for translation ^84,85^. Once ORF1p translocates to the nucleus, it accumulates at the inner NE by directly binding to Lamin B1 in a stress-specific manner. This targeting of the inner NE by ORF1p is paralleled by NE dysmorphology including invaginations and blebs which significantly correlated with ORF1p intensity, both in the nucleus and the inner NE, suggesting that it is the stress-dependent nuclear translocation of ORF1p and the direct binding to Lamin B1 that participate in NE dysmorphology. This conclusion is also supported by the fact that nuclear dysmorphologies under oxidative stress are prevented when ORF1p nuclear import is blocked by nuclear import inhibitors or, through unknown mechanisms, by remodelin.

A direct functional role of ORF1p in NE dysmorphology independent of oxidative stress is further supported by direct ORF1p gain of function experiments showing that neurons over-expressing ORF1p (and not ORF2p) display a significant decrease of nuclear circularity. These experiments also suggest that NE dysmorphology is mediated by the nuclear presence of ORF1p alone and does not rely on the presence of the entire RNP or ORF2p.

NE dysmorphology is associated with NCT deficiencies ^86^ and both have been described to be disturbed in the context of NDs ^52,62,63,87–91^. Indeed, targeting of the NE by ORF1p in neurons was accompanied by NCT deficiencies as revealed by the mislocalization and abnormal and compartmentalized accumulation of several proteins involved in NCT function, some of which interact with ORF1p directly (KPNB1 and NUP153). KPNB1 sequestration by binding to ORF1p might alter nuclear import but could also inhibit its important disaggregase function ^92^. Analogous, the abnormal localization of NUP153 in the nucleoplasm suggests an alteration of its physiological function, potentially mediated by the interaction with ORF1p as illustrated by the accumulation of nucleoplasmic PLA foci. Nucleoporins are long-lived proteins ^93,94^ and rely on cell division for their renewal. Post-mitotic neurons do not divide and nucleoporins therefore are not renewed ^95,96^. Their continuous functioning in non-dividing cells throughout the life time of an individual is thus of crucial importance and any perturbations will have potentially irreversible consequences ^95^. Abnormal intranuclear and cytoplasmic accumulation of certain nucleoporins as we observed for NUP153 in stressed neurons has also been described in the context of other NDs (i.e, HD)^91^. Possible cytoplasmic sequestration of NUP98 by tau has also been suggested to lead to NCT perturbation in AD ^97^. Similarly, delocalization and abnormal cytoplasmic accumulation of RanGAP1 we observed in stressed neurons, have also been described in several ND-affected tissues ^58^. Our results suggest that ORF1p might act as a sponge to alter the activity of certain proteins involved in NCT. Together, our results suggest a possible scenario in which ORF1p might sequester and/or mislocalize NE components such as Lamin B1 or NUP153 (see below).

NCT deficiencies have been consistently linked to the abnormal subcellular distribution of TDP-43, a nuclear protein involved in RNA metabolism ^98,99^, mutated in ALS/FTD ^100–103^ and mislocalized to the cytoplasm in numerous NDs ^58,61,62^ not linked to TDP-43 mutations. Indeed, in our model, NCT deficiencies as a consequence of mislocalized NCT proteins resulted in the cytoplasmic accumulation of TDP-43. TDP-43 exits the nucleus through passive ^104^ or active nuclear export ^105^ through nuclear pores. We also observed an increase in the nuclear pool which could be indicative of a perturbed export mechanism of this shuttling protein ^106^.

NE alteration and NCT defects are emerging pathogenic features in several NDs ^87,97,107^ including HD ^90^, ALS/FTD ^63^, PD ^52^, AD ^97,107^ and more generally in the context of tauopathies ^108^. For example, aggregated Tau directly interacts with the nuclear pore complex^97^ and oligomeric Tau with lamin proteins ^109^ resulting in loss of nuclear integrity including NCT failure ^86^, similar to what we describe here. Loss of nuclear integrity in the form of NE deformation and loss of NE function has been extensively described in the context of aging ^110^ including in the brain ^111^ with associated changes in chromatin organization and NCT dysfunction ^107,111–113^. Progerin, a mutant form of Lamin A, is the cause of HGPS. Similar to mutant Tau, it accumulates at the NE and disrupts NE morphology, NCT and chromatin organization ^69^. In cellular and mouse models of accelerated aging, these alterations can be successfully corrected by treatment with the small molecule remodelin ^69,70^. Remodelin acts by inhibiting the acetyltransferase NAT10 ^69^. In our model of oxidative stress, pre-treatment of neurons with remodelin 24 h prior to arsenite application restored, just as in progeria cells, NE abnormalities. In parallel, remodelin decreased nuclear ORF1p while not affecting cytoplasmic ORF1p content. Remodelin might thus, through the restoration of the nuclear architecture, prevent ORF1p nuclear translocation and therefore further ORF1p-induced NE dysmorphology. Nuclear lamina-alterations do not only affect NCT but alter global chromatin organization ^114^ and potentially cell type specific gene expression ^115^. Chromatin organization in cells is spatial and specific heterochromatin domains are directly linked to the nuclear lamina, called lamina-associated domains (LADs). One marker associated with LADs is H4K20me3, a repressive heterochromatin mark known to silence LINE-1 elements ^57^. Indeed, oxidative stress induced a global loss of H4K20me3 which is rescued by remodelin pre-treatment. Together, these data indicate that remodelin can preserve and/or restore nuclear integrity of neurons including chromatin disorganization induced by oxidative stress possibly or at least partly through an inhibitory effect on ORF1p nuclear translocation. The destructuration of nuclear lamina-anchored heterochromatin as indicated by the delocalization of H4K20me3 from the nuclear periphery could lead to the expression of genes normally repressed in post-mitotic neurons leading to neuronal fate loss, an emerging feature of aging and NDs ^116^. Indeed, mutated Tau can lead to chromatin relaxation, re-expression of certain fetal genes in the brain in a fly model ^18^ and activation of TEs ^14,18^. Further, the loss of H4K20me3 at the nuclear periphery could either be the source of LINE-1 activation (as H4K20me3 is a repressive mark known to decorate repeat DNA ^117,118^) or the consequence of nuclear ORF1p translocation leading to a decrease in H4K20me3 through a yet to be identified mechanism. This would indeed lead to the amplification of the initial stress-induced increase in LINE-1 via a positive feedback loop. Another possibility is an adaptive response aimed at protecting the genome from damage as a short-term response to deformation ^119–121^.

Recent studies have shown that aggregated proteins such as Tau ^83,122^, mutant Huntingtin ^90^ and C9orf72 poly-GA proteins ^58^ can target the NE and alter its integrity and function. Our data suggests that ORF1p might have a pathogenic action similar to ND-linked proteins by perturbing nuclear integrity. In addition, stress-granule independent cytoplasmic TDP-43 droplets can drive nuclear import defects and cell death ^123^. Whether nuclear import defects are driven by a direct ORF1p dependent mechanism or via ORF1p-dependent TDP-43 cytoplasmic accumulation remains to be determined.

Condensation is a feature of ORF1p which is intrinsic to its role in LINE-1 mobilization ^124^. We observed large ORF1p foci at the level of NE invaginations and an increase in the size of cytoplasmic and nuclear ORF1p foci (dots) indicative of ORF1p condensation. Whether ORF1p itself might be aggregation-prone or participate in the organization of LLPS (Liquid-liquid phase separation)-related cytoplasmic or nuclear membraneless organelles requires further investigation.

Altogether, we provide evidence that the LINE-1 encoded protein ORF1p, independent on the LINE-1 RNP or ORF2p, translocates to the nucleus upon cellular stress or overexpression in an importin-beta and NUP153 involving process. Upon translocation, ORF1p binds to Lamin B1 and sequesters and/or mislocalizes this protein, which is essential to NE integrity, leading to further deformation of the NE. In parallel, while entering the nucleus through nuclear pores, ORF1p binds to NUP153, leading to its mislocalization to the nucleoplasm and the cytoplasm and thereby contributing to NCT deficiencies including the cytoplasmic accumulation of TDP-43. Nuclear localization and the correlation of ORF1p nuclear intensity with the loss of nuclear circularity in neurons of PD patients suggest that ORF1p might fulfil a similar pathological function in neurodegenerative disease.

NE deformation is associated with chromatin disorganization as illustrated by HP1 loss at the inner NE and global H4K20me3 depletion in the nucleus. Nuclear deformation and chromatin disorganization upon cell stress were rescued by the small molecule remodelin which might indirectly act as an inhibitor of ORF1p nuclear translocation. This suggests that LINE-1 encoded protein ORF1p, for which no pathogenic function had yet been assigned, might play a pathogenic role by targeting the NE. As LINE-1 activation is a recurring observation in several neurological diseases ^125–131^, in aging ^6^ and more generally upon oxidative stress ^5^ and if this activation includes full-length LINE-1 elements with coding potential, the result could be an increase in ORF1p, in addition to ORF2p and LINE-1 RNA. Passing a certain threshold and in conjunction with a fragilization of the NE either through aging, ND-linked mutated proteins known to affect the NE or oxidative stress, this then might trigger the nuclear translocation of ORF1p contributing to NE deformation and NCT dysfunction. One consequence we identified is the cytoplasmic accumulation of TDP-43 which could favor its aggregation ^132^, one of the hallmarks of several NDs ^58,61,62^.

Activation of LINE-1 elements in neurons, through ORF2p-induced genomic instability ^5^, ORF2p-induced inflammation ^6,8,45^ or LINE-1 RNA mediated heterochromatin erosion ^133^ and, as we show here, through the ORF1p-dependent fragilization of nuclear integrity could contribute in multiple ways to pathological features common to several NDs. This reinforces the idea that LINE-1 elements might represent a novel therapeutic target for neuroprotection.

## Acknowledgements

RZ acknowledges the Fondation Recherche Alzheimer (FRA) for a Ph.D. fellowship and is enrolled with the Ecole Doctorale ED3C at Sorbonne University. We are very grateful to Raphaël Rodriguez and Ludovic Colombeau for having synthetized and provided the small molecule remodelin. This work was supported by grants to JF from the Fondation de France (00086320), the Fondation du Collège de France, the Fondation NRJ/Institut de France, the National French Agency for Research (ANR-20-CE16-0022 NEURAGE) and the Fondation Alzheimer. We gratefully acknowledge Julien Dumont and the Collège de France Orion imaging facility (IMACHEM-IBiSA), member of the French National Research Infrastructure France-BioImaging (ANR-10-INBS-04), which received support from the program«Investissements d’Avenir» ANR-10-LABX-54 MEMOLIFE. We also thank the Fondation Bettencourt Schueller for their support. Human post-mortem brain material and associated data were obtained from the “Brainbank Neuro-CEB Neuropathology Network” which is supported by ARSLA, CSC, France DFT, Fondation ARSEP, Fondation Vaincre Alzheimer, France Parkinson.

## Author contributions

RZ performed most of the experiments. OMB contributed experimentally, PM and HM wrote the scripts for the automated image analysis workflow, RLJ and JF supervised the study and wrote the paper with RZ. JF received the funding.

