## Supplementary Figure Legends for "Nuclear translocation of LINE-1 encoded ORF1p alters nuclear envelope integrity and disrupts nucleocytoplasmic transport in human neurons"

**Suppl. Figure 1. Differentiated LUHMES cells express neuronal and dopaminergic markers TuJ1 and TH.**

(A) Experimental protocol of the differentiation of neuronal precursor cells (LUHMES) into mature, post-mitotic dopaminergic neurons in 5 days. (B) Quantification of the fluorescence intensity of TH and TuJ1. n > 100 neurons were quantified per condition, a representative experiment of 3 independent experiments (3 wells per condition), mean ± SEM, One-way ANOVA with Tukey’s multiple comparisons test. (C) Analysis of TH protein levels by Western blot. (D) Immunofluorescence of TuJ1 and TH in the absence and the presence of oxidative stress. (E) Arsenite treatment does not affect the neuronal and the dopaminergic phenotypes of differentiated LUHMES cells. n > 100 neurons were quantified per condition, a representative experiment of 3 independent experiments (3 wells per condition), mean ± SEM, two-tailed t test. (F) LINE-1 RNA levels were increased as shown by RT-qPCR using primers targeting different LINE-1 regions. Cycle thresholds were obtained from 3 wells per conditions from 2 independent experiments, using the ddCt method relative to the expression of reference genes HPRT and TBP. Data are represented as the mean ± SEM, two-way ANOVA with Tukey’s multiple comparisons test (*p < 0.5, **p < 0.01, ***p < 0.001, ****p < 0.0001). Scale bar, 5 μm.

**Suppl. Figure 2. Increased DNA damage in differentiated LUHMES cells following oxidative stress is restored upon stavudine treatment.**

(A) ssDNA Immunostaining in arsenite treated LUHMES cells. n ≥ 50 neurons were quantified per condition, a representative experiment of 2 independent experiments (3 wells per condition). (B) Nuclear deformations are paralleled by an increase of Lamin B1 at the inner NE and in the nucleoplasm, mean ± SEM, two-tailed t test. (C, D) Quantification of ORF1p nuclear intensity and the nuclear circularity in the presence or the absence of import inhibitor drug ivermectin (IV) only or importazole (IMP) only in stressed and control cells. n > 100 neurons were quantified per condition, a representative experiment (3 wells per condition) of 2 independent experiments, mean ± SEM, One-way ANOVA with Tukey’s multiple comparisons test. (*p < 0.5, **p < 0.01, ***p < 0.001, ****p < 0.0001). Scale bar, 5 μm.

**Suppl. Figure 3. Nuclear dysmorphologies, and abnormal ORF1p increase in differentiated LUHMES cells following oxidative stress are not restored upon stavudine treatment.**

(A) Immunostaining of Lamin B1 following arsenite treatment in the presence or the absence of stavudine and quantification of the circularity index. (B) Co-Immunostaining of ORF1p and Lamin B1 following arsenite treatment in the presence or the absence of stavudine and quantification of Lamin B1 in the nucleoplasm. (C) Quantification of ORF1p intensity both in the cytoplasm and the nucleus following stress in the presence or the absence of stavudine. n > 100 neurons were quantified per condition (3 wells per condition) from 1 experiment, mean ± SEM, two-tailed t test (*p < 0.5, **p < 0.01, ***p < 0.001, ****p < 0.0001). Scale bar, 5 μm.
