## Supplementary Figures for "Nuclear translocation of LINE-1 encoded ORF1p alters nuclear envelope integrity and disrupts nucleocytoplasmic transport in human neurons"

### Slide 1
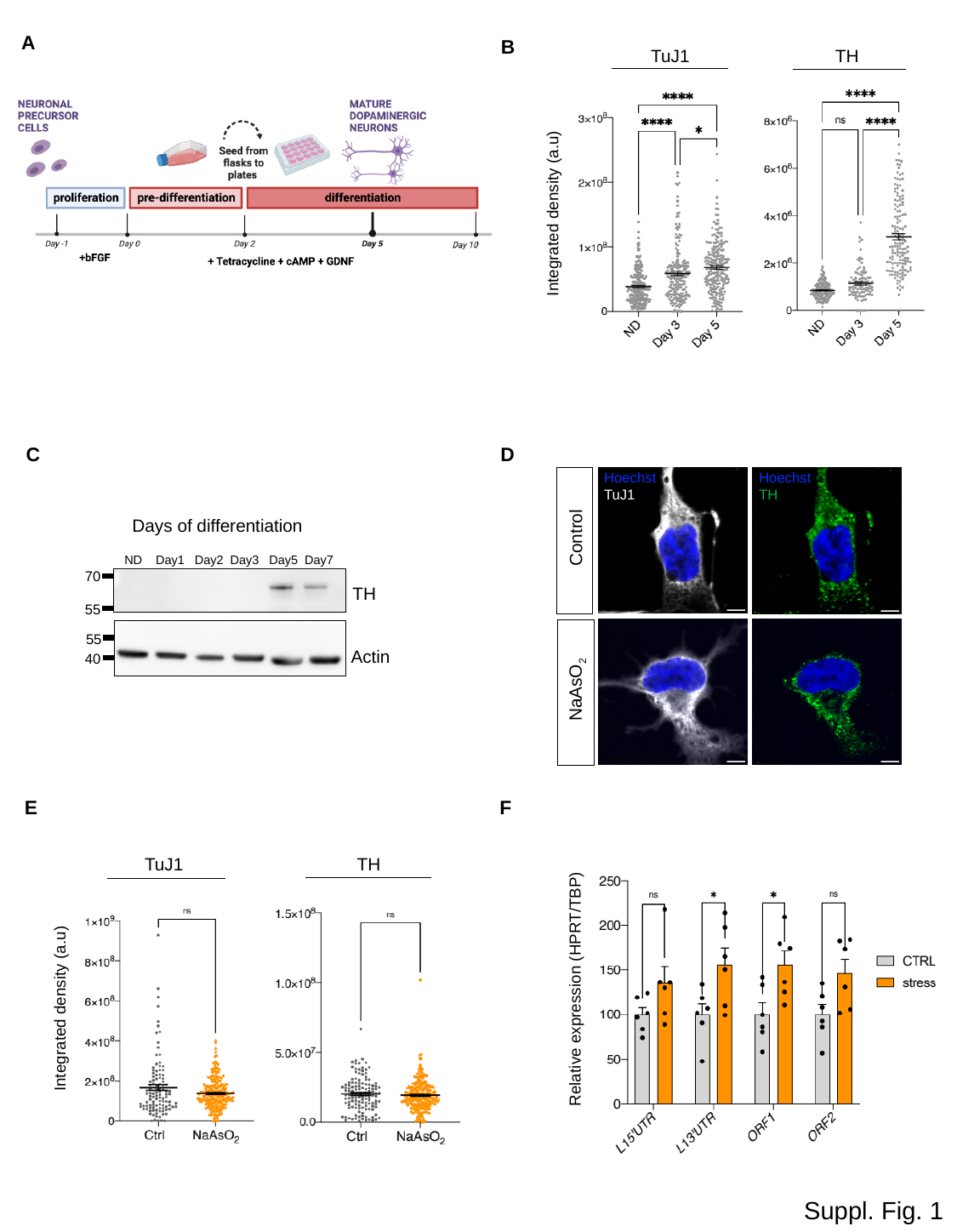

A
B
TuJ1
TH
 Integrated density (a.u)
D
C
Hoechst
TuJ1
Hoechst
TH
 Control
 NaAsO_2_
Days of differentiation
 ND Day1 Day2 Day3 Day5 Day7
70
TH
55
55
Actin
40
E
F
TuJ1
TH
 Integrated density (a.u)
Relative expression (HPRT/TBP)
Suppl. Fig. 1

### Slide 2
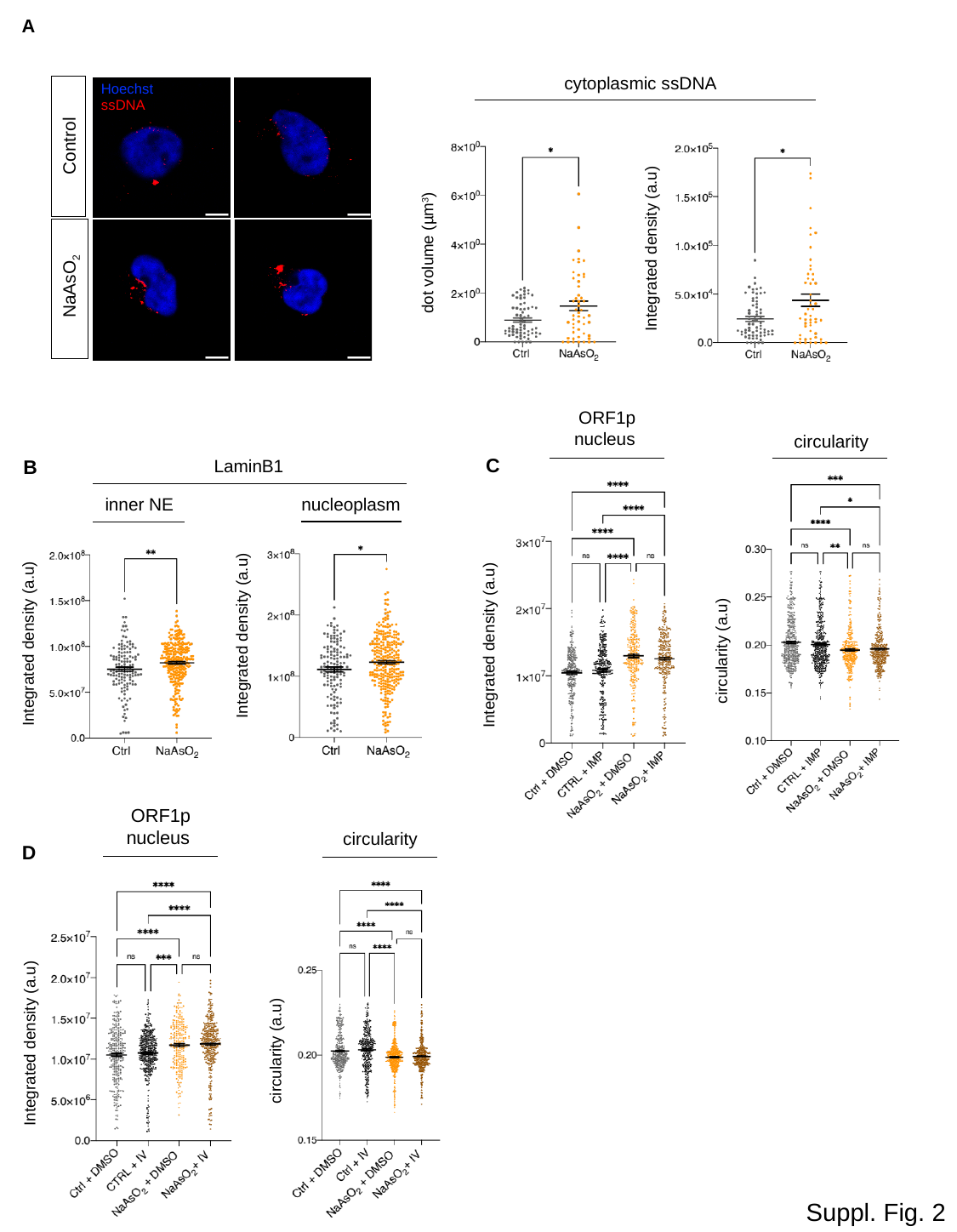

A
cytoplasmic ssDNA
Hoechst
ssDNA
Control
 Integrated density (a.u)
 dot volume (µm^3^)
 NaAsO_2_
ORF1p
nucleus
circularity
C
B
LaminB1
inner NE
nucleoplasm
 Integrated density (a.u)
 Integrated density (a.u)
circularity (a.u)
 Integrated density (a.u)
ORF1p
nucleus
circularity
D
circularity (a.u)
 Integrated density (a.u)
Suppl. Fig. 2

### Slide 3
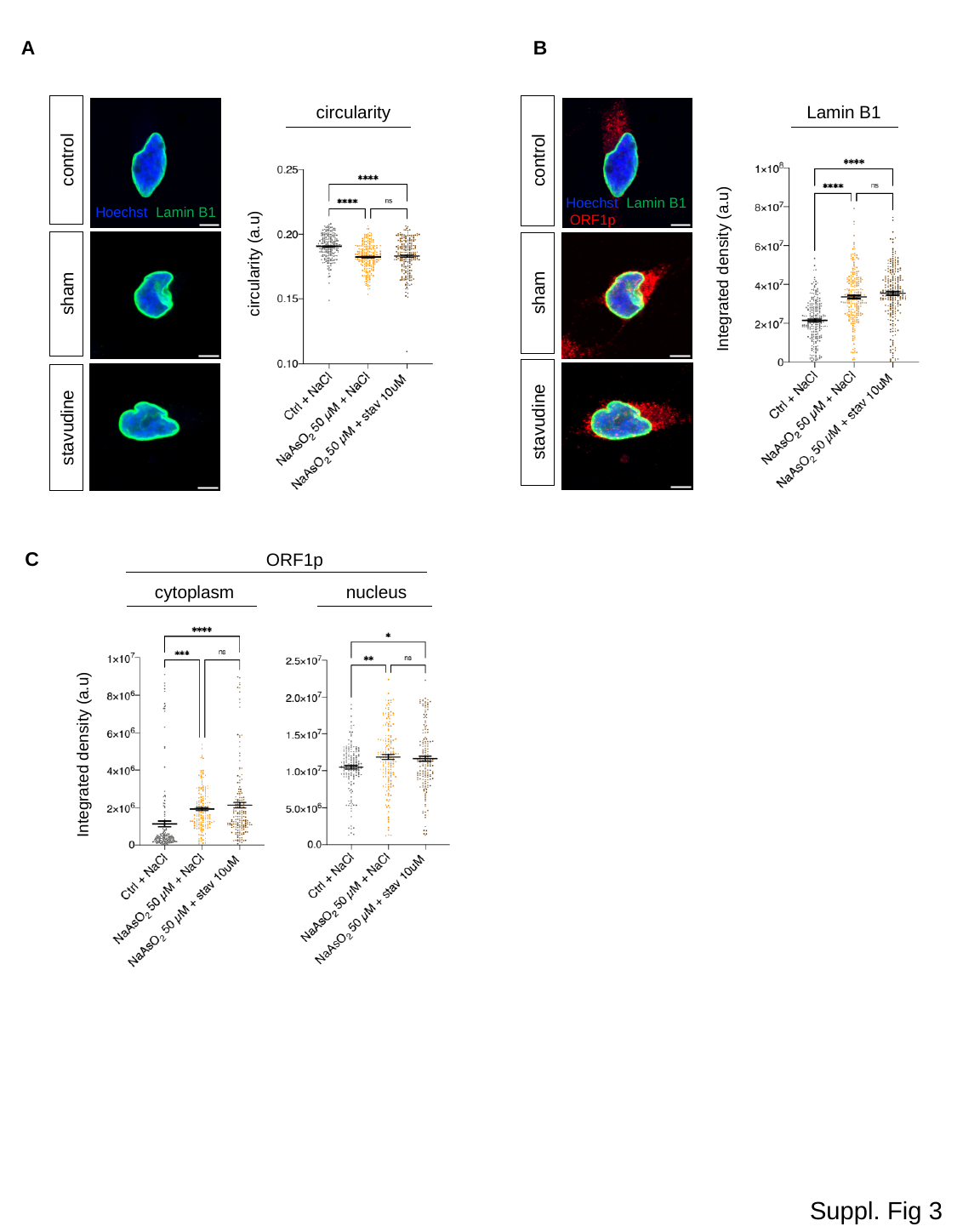

A
B
circularity
Lamin B1
control
control
Hoechst Lamin B1 ORF1p
Hoechst Lamin B1
circularity (a.u)
 Integrated density (a.u)
sham
sham
stavudine
stavudine
C
ORF1p
cytoplasm
nucleus
 Integrated density (a.u)
Suppl. Fig 3
